# Chromosomal integration of *bla*_CTX-M_ redefines the evolutionary trajectory of antibiotic resistance in *E. coli* through lineage-structured expansion

**DOI:** 10.64898/2026.10.07.757360

**Authors:** Maxime Danjean, Thibaut Morel-Journel, Valentine Berti, Yasmine Benhadid-Brahmi, Signara Gueye, Houéfa Christelle Lougbegnon, Hervé Jacquier, Erick Denamur, Stéphane Bonacorsi, Patricia Siguier, François Cornet, Manuel Campos, Marie Petitjean, André Birgy, Laurence Armand-Lefevre

**Affiliations:** Université Sorbonne Paris Nord et Université Paris Cité, INSERM, IAME, Paris, France; Service de Microbiologie, Hôpital Robert-Debré, AP-HP, Paris, France; Assistance Publique-Hôpitaux de Paris (AP-HP), Hôpital Bichat-Claude Bernard, Laboratoire de Bactériologie, Paris, France; Bacteriology Unit, Department of Prevention, Diagnosis, and Treatment of Infections, Henri-Mondor University Hospital, AP-HP, Creteil, 94000, France; INSERM U955 (IMRB), University of Paris-Est, Créteil, 94000, France; Laboratoire de Microbiologie et Génétique Moléculaires, Centre de Biologie Intégrative, Université de Toulouse, CNRS, UPS, 165 Rue Marianne Grunberg-Manago, 31400 Toulouse, France

**Keywords:** Extended-spectrum beta-lactamase (ESBL), *Escherichia coli*, CTX-M, chromosomal integration, evolution

## Abstract

The global CTX-M pandemic is considered a plasmid-driven phenomenon, yet chromosomal integration of *bla*_CTX-M_ has been reported in various settings. Screening 451,174 publicly available *E. coli* genomes and retaining 2,269 CTX-M-*E. coli* long-read or hybrid assemblies, we found that 33.5% carried at least one chromosomally integrated *bla*_CTX-M_. Chromosomal integration was strongly structured by *E. coli* lineage and CTX-M variant, reaching 68.5% in ST38 and 43.9% in ST131. In the latter, integration rates varied between subclades with modelling, suggesting recent acceleration within the ST131-C2 subclade. Additional resistance genes were frequently co-integrated with *bla*_CTX-M_ (33%), with architectures largely inherited from plasmid-borne structures. Integrations were mainly IS*Ecp1*- or IS*26*-mediated (77.3% and 18.9%, respectively) and occurred across the entire *E. coli* chromosome without spared zones. *bla*_CTX-M_ chromosomal integration is a widespread, lineage-structured component of the CTX-M pandemic, providing stable vertical inheritance that complements or may supplant plasmid-mediated dissemination in successful ESBL-*E. coli* clones.

## INTRODUCTION

Antimicrobial resistance (AMR) is a major global health threat. In 2021, it was estimated that approximately 5 million deaths worldwide were associated with infections caused by resistant bacteria, including 1.14 million directly attributable to AMR^1^. *Escherichia coli* ranks among the leading causes of AMR-associated deaths while simultaneously thriving as a commensal of the human and animal gut microbiota^1^.

Among resistant *E. coli*, extended-spectrum beta-lactamase-producing *E. coli* (ESBL-*E. coli*) have emerged as one of the most widespread multidrug-resistant bacteria globally and are classified by the World Health Organization as critical priority pathogens requiring urgent research and development of new therapeutics^2^.Their global expansion has been largely driven by CTX-M-type enzymes, particularly CTX-M-15, which have disseminated since the 1990s across humans, animals, and the environment^3–5^. This expansion has led to a steady increase in the prevalence of ESBL-producing Enterobacterales, with global human carriage estimated at 10-30% between 2014 and 2024 with substantial geographic variation^6^. The pandemic success of CTX-M-producing *E. coli* is likely multifactorial^6,7^, resulting from both the efficient horizontal transfer of *bla*_CTX-M_ genes and the global dissemination of highly adapted clones. Notably, the *E. coli* Sequence Type 131 (ST131) lineage has played a pivotal role through acquisition of *bla*_CTX-M-15_-carrying plasmids, combining multidrug resistance with high colonization and transmission capacity^6,8^. Today, ST131 remains the dominant ESBL-producing lineage worldwide, alongside emerging clones such as ST1193^9^, ST10, ST405, ST38^8^ and more recently ST69^10^, across both high, low- and middle-income countries. ESBL-*E. coli* populations are more diverse in low- and middle-income countries, where higher prevalence and stronger human-animal-environment connectivity may broaden transmission pathways^4^. The *bla*_CTX-M_ genes originate from the chromosome of the environmental Enterobacterales genus *Kluyvera,* where they function as intrinsic beta-lactamase genes^11^. Their mobilization was enabled by insertion sequences (IS) (e.g., IS*Ecp1*), which facilitated their capture and integration into mobile genetic elements (MGEs) such as integrons and transposons^12^. These elements, frequently located on conjugative multidrug-resistance plasmids, have driven efficient horizontal gene transfer across bacterial species and cross-sector dissemination under antibiotic selection pressure^4,7^.Consequently, acquired beta-lactam resistance in Enterobacterales has long been considered primarily plasmid-borne, with horizontal transfer widely driving their dissemination, a paradigm that shaped our understanding of AMR dynamics^13^. Plasmids were viewed both as a major threat, due to their conjugative transfer and capacity for horizontal dissemination^13,14^ and a potential vulnerability, as their biological cost could lead to plasmid loss in the absence of antibiotic pressure^15,16^. However, accumulating evidence challenges this model. Increasing reports indicate chromosomal integration of *bla*_CTX-M_ and other antimicrobial resistance genes (ARGs) in *E. coli*^17–21^. Consistently, data from Madagascar and Niger show predominant chromosomal integration of *bla*_CTX-M_ (57% and 50% respectively) across sectors, often associated with co-integrated ARGs^22,23^. This chromosomal location may reduce plasmid-associated fitness costs^15,24^ and favor vertical inheritance of resistance determinants, potentially contributing to their persistence in bacterial populations. Chromosomal integration may therefore represent a significant route toward long-term stabilization of acquired resistance, with potential epidemiological consequences.

Precise identification of the genetic support of CTX-M resistance has been enabled by advances in whole-genome sequencing (WGS) and *in silico* plasmid identification tools^25,26^. While short-read WGS provides high-resolution insights into clonal dissemination and genomic content^27^, it remains limited in resolving plasmid architecture and chromosomal integration events, owing to repetitive sequences and conserved mobile element backbones^28,29^. *In silico* tools inferring *bla*_CTX-M_ localization from genomic context complement short-read approaches but remain limited by assembly fragmentation and are blind to mosaic chromosomal insertions of plasmid origin, potentially underestimating integration events^25^. In contrast, long-read sequencing and hybrid-assembly approaches allow contiguous genome reconstruction^29^, including complex plasmids and chromosomes, thereby improving detection of *bla*_CTX-M_ chromosomal integration. Although increasingly adopted due to technical improvements and declining costs, these technologies remain less automated than short-read sequencing and are not yet widely accessible, limiting their large-scale implementation. As a result, the respective contributions of chromosomal and plasmid supports, as well as the precise epidemiology of *bla*_CTX-M_ chromosomal integration in *E. coli*, remain poorly characterized. To address this gap, we sought to assess the prevalence and epidemiology of chromosomal *bla*_CTX-M_ integration using a large-scale collection of long-read or hybrid assemblies. We further characterized the distribution of *bla*_CTX-M_ variants and the molecular epidemiology of isolates harboring chromosomal integration events. By clarifying the evolutionary and epidemiological dynamics of chromosomal integration, this work provides insight into a potential transition from predominantly mobile to genomically stabilized resistance, with implications for the future trajectory and control of AMR.

## RESULTS

### A long-read genome collection reveals widespread chromosomal *bla*_CTX-M_ integration in ESBL-producing *E. coli*

To quantify chromosomal *bla*_CTX-M_ integration at scale, we screened all 451,174 *E. coli* available genomes in the NCBI database and retained long-read or hybrid assemblies meeting predefined quality criteria. Among them, 6,873 met quality criteria of which 2,507 harbored at least one *bla*_CTX-M_ gene from 666 BioProjects. After removal of closely related duplicates based on BioProject origin, cgMLST distance (<10 allele differences) and identical resistome, this yielded a non-redundant collection of 2,269 genomes harboring 2,676 *bla*_CTX-M_ genes (**Extended Data Fig. 1**) (https://doi.org/10.5281/zenodo.20934804).

The population structure genome collection reflects the known epidemiology of ESBL-*E. coli,* with the ST131 as the most represented lineage (17.9%; 407/2269), followed by ST10 (5.5%; 124/2269), ST38 (5.4%; 123/2269), ST410 (4.9%; 112/2269), ST167 (4.7%; 106/2269) and ST405 (3.5%; 80/2269). The *bla*_CTX-M_ repertoire was dominated by CTX-M-15 (40.8%; 1093/2676), CTX-M-55 (15.7%; 421/2676), CTX-M-14 (15.2%; 407/2676) and CTX-M-27 (10.1%; 271/2676) (**Fig. 1a-b**).

**Figure 1.**
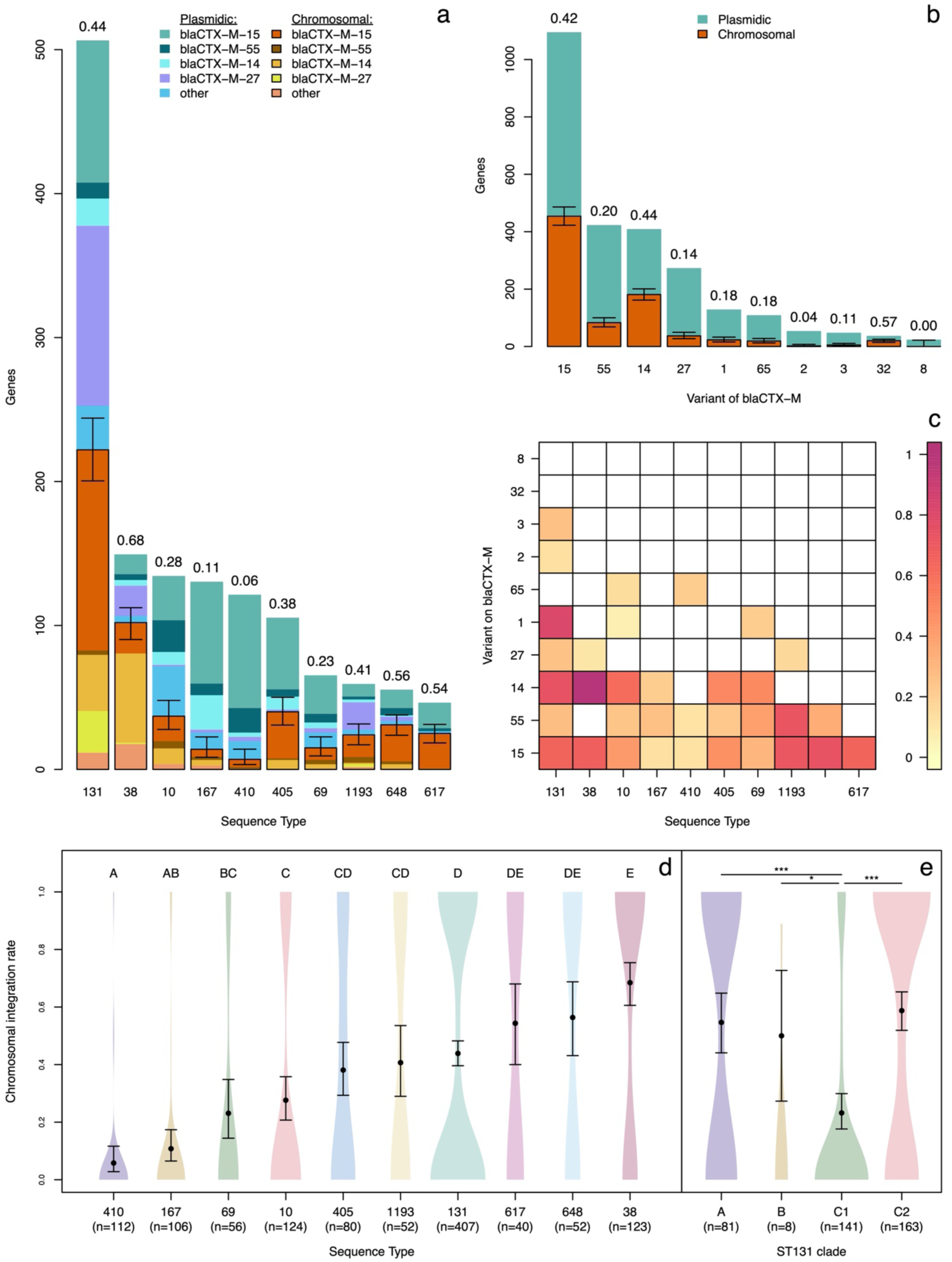
Prevalence of chromosomal *bla*_CTX-M_ gene integration across ESBL-producing *Escherichia coli* Sequence Types, subclades and variants. **a,** Number of *bla*_CTX-M_ genes and variants located on the chromosome or plasmid among the ten main *E. coli* Sequence Types (STs) **b,** Number of *bla*_CTX-M_ genes located on the chromosome or plasmid among the ten most frequent *bla*_CTX-M_ variants. In **a-b**, total bar height corresponds to the cumulative number of *bla*_CTX-M_ genes identified in each category, irrespective of genomic location. Values displayed above each bar indicate the proportion of chromosomally integrated *bla*_CTX-M_ genes within the corresponding category. Error bars represent the 95% confidence interval of the estimated proportion of chromosomal *bla*_CTX-M_ genes. **c,** Heatmap of the integration rate for given couples of variant and ST. Only integration rates for couples observed at least five times are represented. **d-e,** Distribution of chromosomal integration rates across ESBL-Ec Sequence Types and across subclades of ST131. Violin plots (color) representing the distribution of observed integration rates for the 10 main STs (**d**) and for subclades of ST131 (**e**), with the estimates from the logistic regressions with their 95% confidence intervals (black). The number of strains considered is in parentheses. Significant differences are computed according to Tuckey’s HSD tests. **d,** pairwise differences were summarized through compact letter display: Sequence Types sharing a letter (from A to E) do not differ significantly (Tuckey’s HSD test p > 0.05). **e,** significant differences between subclades are noted, with “*” for a p-value lower than 0.05 and “***” for a p-value lower than 0.001. In **a-e,** chromosomal and plasmid locations were inferred from genome assembly analyses.

Overall, we first observed that 32.3% (864/2,676) of the *bla*_CTX-M_ genes, were located on the chromosome and 67.7% (1,812/2,676) on plasmids, corresponding to chromosomal integration in 33.5% (761/2,269) of genomes, establishing that *bla*_CTX-M_ chromosomal integration is a frequent feature of ESBL-*E. coli* (**Fig. 1b**).

### Chromosomal *bla*_CTX-M_ integration is structured by lineage rather than randomly distributed across the *E. coli* population

To determine whether chromosomal integration of *bla*_CTX-M_ was shaped by specific CTX-M variants or *E. coli* genetic background, we analyzed its distribution across CTX-M variants, phylogroups, and Sequence Types.

Integration rates varied markedly across the four CTX-M variants: 13.7% (37/271) for CTX-M-27, 19.7% (83/421) for CTX-M-55, 41.5% (454/1,093) for CTX-M-15, and 45.4% (181/407) for CTX-M-14 (**Fig. 1b**).

*E. coli* core-genome phylogenetics revealed that genomes carrying chromosomally integrated *bla*_CTX-M_ genes tended to cluster within specific phylogenetic backgrounds rather than being scattered across the tree (**Fig. 2**). Chromosomal integration was significantly more prevalent in the extraintestinal-associated B2 and D phylogroups (46.8%; 275/587 and 47.1%, 162/344, respectively) than in the commensal-associated A and B1 phylogroups (26.5%, 168/635 and 21.4%, 78/365) (p<0.001) (**Extended Data Fig. 2**). Among STs, differences in *bla*_CTX-M_ chromosomal integration rates were even more pronounced, spanning a more than ten-fold range, from 5.8% (7/121) in ST410 to 68.5% (102/149) in ST38 (**Fig. 1a,d; Extended Data Fig.S3. Supplementary Table S1; Supplementary Table S2**). Other major epidemic ESBL-*E. coli* clones also showed markedly variable rates: 56.4% (31/55) in ST648, 43.9% (222/506) in ST131, 40.7% (24/59) in ST1193 and 27.6% (37/134) in ST10 (**Fig. 1a,d; Supplementary Table S1**).

**Figure 2.**
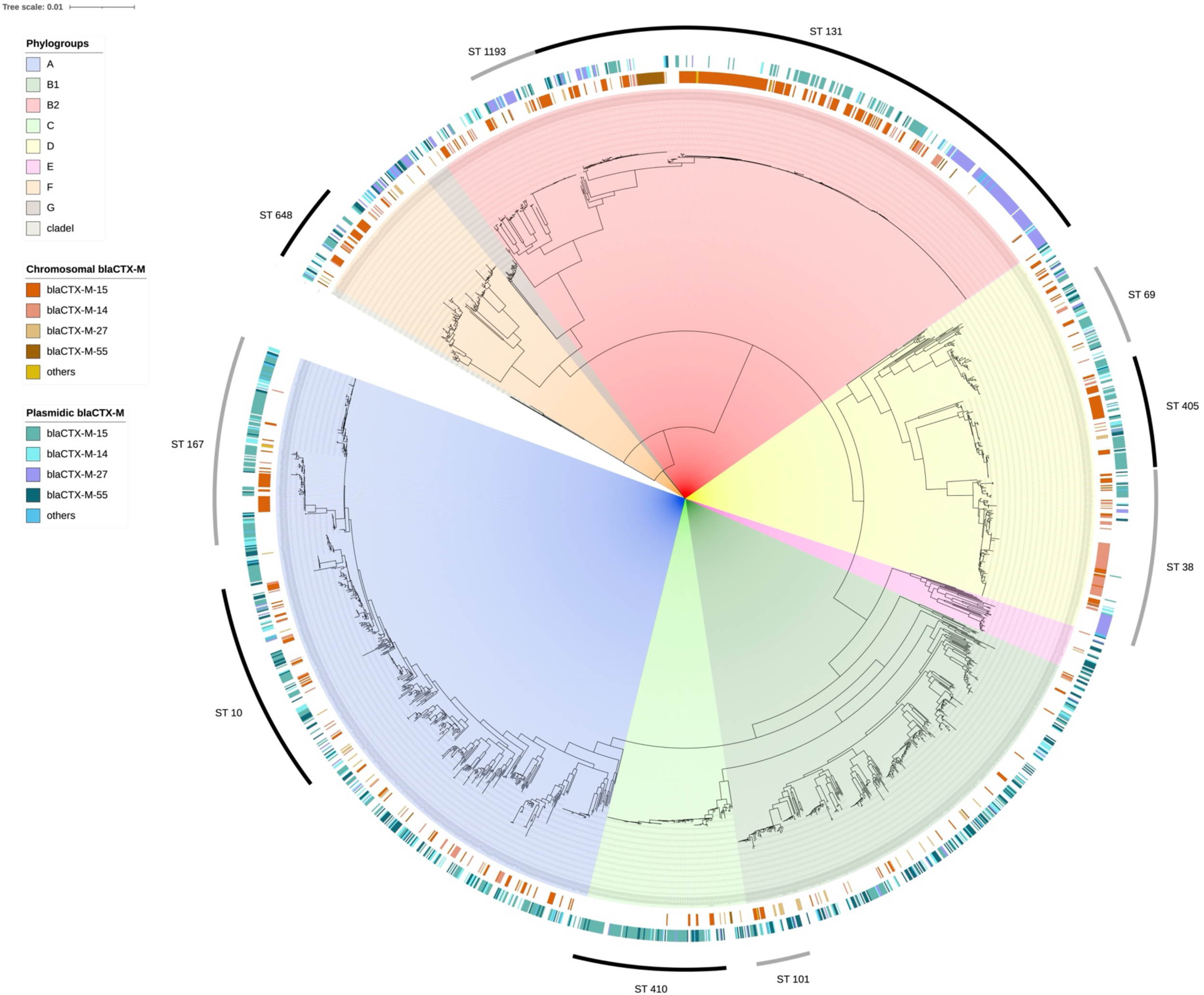
Maximum-likelihood phylogeny reconstructed from the persistent genome partition of the 2,269-genome *E. coli* pangenome using a GTR+G model and rooted with an *Escherichia fergusonii* outgroup (genome accession: GCA_020097475.1). Branches are highlighted depending on the phylogenetic group of the *E. coli* genome. The inner ring represents the *bla*_CTX-M_ variants located on the chromosome, ring two indicates the *bla*_CTX-M_ variants located on plasmids, ring three indicates the ten main sequence types (STs).

Chromosomal integration rates also varied within each ST according to the CTX-M variant, highlighting ST/variant-specific combinations: in ST131, integration reached 67.2% (39/58) for CTX-M-14 versus 18.8% (29/154) for CTX-M-27 and in ST38, it reached 93.9% (62/66) for CTX-M-14 versus 4.5% (1/22) for CTX-M-27 (**Fig. 1c; Extended Data Fig. 3**).

Overall, chromosomal integration rates were strongly structured by both CTX-M variant and host genetic background. Specific ST/variant combinations, consistently over-represented, reflect either a genuine propensity for integration or early integration followed by clonal expansion.

### Chromosomal *bla*_CTX-M_ integration is largely independent of carbapenemase co-production across *E. coli* lineages

As a subset of the collection, 671 ESBL-*E. coli* genomes carried at least one carbapenemase gene. We assessed whether carbapenemase gene co-occurrence influenced chromosomal *bla*_CTX-M_ integration. Overall, *bla*_CTX-M_ integration rates did not differ significantly between ESBL-only and ESBL+carbapenemase-*E. coli* genomes (34.1%; 545/1,598 vs 32.2%; 216/671 respectively; *p* = 0.408) (**Supplementary Fig. S1**). We then examined two carbapenemase-associated high-risk clones, ST167 and ST410. In ST167, *bla*_CTX-M_ chromosomal integration rates were comparable between ESBL-only and ESBL+carbapenemase-*E. coli* (16.7%, 3/18 vs 8.0%, 7/88, respectively; *p* = 0.368). In ST410, however, *bla*_CTX-M_ chromosomal integration was significantly more prevalent in ESBL-only than in ESBL+carbapenemase-*E. coli* (17.2%, 5/29 vs. 2.4%, 2/83; *p* = 0.012) (**Supplementary Fig. S1**).

Overall, carbapenemase gene co-occurrence was not associated with difference in chromosomal *bla*_CTX-M_ integration, but the association was observed within ST410 points to a lineage-specific effect.

### ST131 subclades are consistent with distinct accumulation dynamics of chromosomal *bla*_CTX-M_

To refine our analysis, we then examined integration dynamics at the intra-ST level. ST131 offered a unique opportunity to examine intra-ST integration dynamics combining the largest sample size (407 genomes), frequent chromosomal integration (43.9%), and a well characterized phylogenetic structure comprising four subclades (A, B, C1 and C2) with documented divergence dates (A/B divergence ca. 1890, B/C ca. 1990, C1/C2 ca. 2000). The four subclades exhibited substantial differences in chromosomal integration rates (**Fig. 1e**), most strikingly between C1 (23.2%) and C2 (58.7%) subclades, despite the recency of their divergence. We modelled the trajectories of chromosomal integration along the ST131 phylogeny to assess subclade-specific dynamics of their accumulation (**Supplementary Note 1**). Based on the phylogenetic structure of ST131 subclades, we inferred branch-specific accumulation speeds resulting in the current differences observed. The model indicates that, while similar accumulation dynamics were estimated for the ancestral lineages A and B, accumulation slowed in the C1 subclade after its divergence from B and accelerated markedly in C2 after its divergence from C1. The reconstructed trajectories suggest that C2 has recently overtaken all other ST131 subclades, reaching the highest predicted proportion of chromosomally integrated *bla*_CTX-M_ genes, driven by a particularly high accumulation speed. (**Extended Data Fig. 4**) (**Supplementary Tables S2-a**, **S2-b, and S3**).

Overall, ST131 subclades showed markedly different chromosomal *bla*_CTX-M_ integration patterns, with exploratory modelling consistent with a more recent increase in C2.

### Chromosomal *bla*_CTX-M_ integration can involve multiple copies or multiple integration within the same chromosome

We next investigated the occurrence of multiple chromosomal *bla*_CTX-M_ integration within individual genomes. Among the 761 genomes carrying at least one chromosomal insertion, 78 (10.2%) harbored multicopy or multi-integration with copy numbers ranging from 2 to 8, most involving identical *bla*_CTX-M_ variants (**Supplementary Table S2-a**). Mixed-variant combinations occurred in only 6 genomes and included CTX-M-15/CTX-M-14, CTX-M-14/CTX-M-55, and CTX-M-172/CTX-M-183 variants.

Multicopy prevalence varied by *bla*_CTX-M_ variant: 17.5% (25/143) for *bla*_CTX-M-14_, 12.2% (9/74) for *bla*_CTX-M-55_, 6.9% (29/420) for *bla*_CTX-M-15_ and 2.8% (1/36) for *bla*_CTX-M-27_. It also differed across STs: 19% (16/84) for ST38, 11.4% (21/184) for ST131, 11.1% (4/36) for ST405, 5.8% (2/35) for ST10, and 3.3% (1/30) for ST648. Stratification by *bla*_CTX-M_ variant within STs revealed that multicopy was predominantly driven by specific ST/variant combinations, most notably *bla*_CTX-M-14_ in ST131 (39.1%, 9/23) and ST38 (22.4%, 11/49), mirroring the ST/variant-dependent pattern observed for chromosomal integration rates overall.

Multicopy *bla*_CTX-M_ loci, observed across unrelated lineages, reflect either repeated independent insertion events or subsequent amplification of integrated elements, supporting a broader process of chromosomal accretion and stabilization of resistance determinants.

### Chromosomal *bla*_CTX-M_ genes are embedded within variant-specific resistance gene islands

As chromosomal integration is believed to have originated from multi-drug resistance plasmids, we explored the associated ARGs surrounding the integrated *bla*_CTX-M_ genes. To characterize the ARG content of chromosomal *bla*_CTX-M_ insertion regions, we defined variant-specific minimal windows using pairwise patristic distances between unrelated genomes, capturing shared insertion-associated sequences while excluding unrelated chromosomal background. (**Supplementary Note 2, Supplementary Table S4**). The resulting windows spanned −16 to +33kb for *bla*_CTX-M-15_, −24 to +5kb for *bla*_CTX-M-55_, −9 to +14kb for *bla*_CTX-M-14_, and −15 to +21kb for *bla*_CTX-M-27_ (**Supplementary Fig. S2; Extended Data Fig. 5**). Overall, 33.6% (256/761) of genomes with chromosomal *bla*_CTX-M_ gene carried at least one additional ARG, in the surrounding region, amounting to 753 associated ARGs (53 distinct ARGs). Among them, the most frequently co-localized genes were *bla*_OXA-1_ and *aac*(6’)-Ib-cr (19.8%; 149/753 each), *aac*(3)-IIa (15.3%; 115/753), *qnr*S1 (8.4%; 63/753) and *tet*(A) (6.1%; 46/753). We identified 73 distinct ARG combinations (“barcodes”) across 256 genomes (259 occurrences) including three only shared between *bla*_CTX-M_ variants. (**Fig. 3a**). *bla*_CTX-M-15_ dominated the diversity, with 48 barcodes and 84.5% (219/259) of all occurrences. The three most frequent barcodes were [*aac*(3)-IIa]_[*bla*_OXA-1_]_[*aac*(6’)-Ib-cr] (21.6%; 56/259), [*qnr*S1] (13.1%; 34/259), [*aac*(3)-IIa]_[*bla*_OXA-1_]_[*aac(6’)-*Ib-cr]_[*tet*(A)] (6.6%; 17/259), resulting in co-chromosomal stabilization of beta-lactams and multiple other resistance determinants (**Fig. 3b-c**., **Supplementary Table S2-c**).

**Figure 3.**
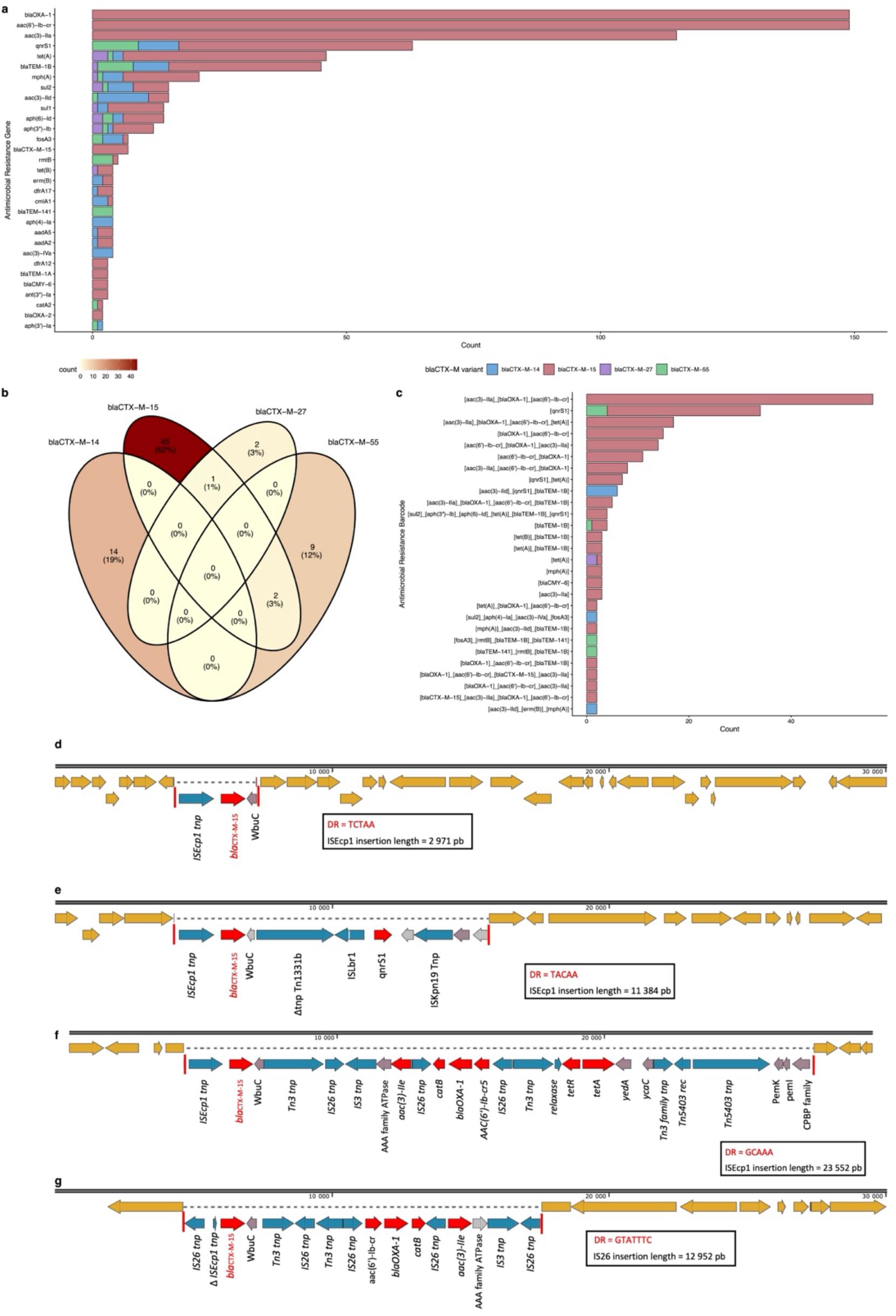
Antimicrobial resistance genes, resistance gene barcodes and mobile-element architectures associated with the main chromosomal *bla*_CTX-M_ variants. **a,** Distribution of antimicrobial resistance genes (ARGs) within variant-specific genomic windows surrounding chromosomal *bla*_CTX-M_ genes. **b,** Venn diagram showing the overlap of the 73 ARGs barcodes among the four main chromosomal *bla*_CTX-M_ variants. Each barcode corresponds to a unique combination of ARGs identified within the variant-specific genomic windows. Numbers indicate barcode counts and percentages. The color gradient reflects the number of barcodes in each region. **c.,** Distribution of ARG barcodes identified around chromosomal *bla*_CTX-M_ genes. Barcodes were defined as the ordered sequence of ARGs within genomic windows surrounding chromosomal *bla*_CTX-M_ genes and were displayed according to the orientation of the *bla*_CTX-M_ open reading frame. **a., c.**, Bars represent the number of occurrences of each ARG or barcode and are colored according to the associated *bla*_CTX-M_.variant. Only ARGs and barcodes detected more than once are shown. All antimicrobial resistance genes barcodes are described in **Supplementary Table S2-c**. **d–g**, Representative chromosomal *bla*_CTX-M-15_ contexts and associated mobile-element architectures. Chromosomal *E. coli* genes are shown in orange, ARGs in red, transposases in blue, hypothetical proteins in grey, and other genes in purple. Red vertical bars indicate target-site duplications (DR, direct repeats). **d**, GCA_013694325.1: 2,971-bp ISEcp1-mediated insertion between *smr*B and *prm*B, flanked by a 5-bp DR (TCTAA). **e**, GCA_014160155.1: 11,384-bp ISEcp1-mediated insertion within *ynb*D, flanked by a 5-bp DR (TACAA). **f**, GCA_050193975.1: 23,552-bp ISEcp1-mediated insertion carrying additional ARGs and disrupting *ydh*S, flanked by a 5-bp DR (GCAAA). **g**, GCA_015138575.1: 12, 952-bp IS26-associated insertion carrying additional ARGs, flanked by a 7-bp DR (GTATTTC).

To assess whether chromosomal ARGs could originate from plasmid-borne structures, we compared chromosomal and plasmidic ARGs and barcodes. Overall, 45.4% (1,029/2,269) of ESBL-*E. coli* genomes carried plasmids harboring ARGs. **(Supplementary Table S2-d; Extended Data Fig. 6a-c).** Most chromosomal ARGs were also detected on plasmids (45/53, 84.9%), representing 98.5% (742/753) of chromosomal ARG occurrences, whereas 43 plasmidic ARGs were never identified on the chromosome. At the barcode level, 41.1% (30/73) of distinct chromosomal barcodes had a plasmidic counterpart, accounting for 71.8% (186/259) of chromosomal barcode occurrences. However, plasmidic ARG frequency did not always predict chromosomal integration: 33.0% (149/452) of *bla*_OXA-1_ and 33.6% (149/443) of *aac*(6’)-Ib-cr were co-integrated versus only 4.8% (21/433) of *mph*(A) and 3.3%14/419) of *sul1*, despite comparable overall frequencies (**Extended Data Fig. 6a-c; Supplementary Fig. S3**.

Chromosomal ARG regions appear largely inherited from plasmid-borne structures, yet their integration is selective and shaped by the surrounding genetic architecture rather than by ARG frequency alone.

### Representative IS*Ecp1*- and IS*26*-associated structures support distinct routes of chromosomal *bla*_CTX-M_ integration (222 words)

To investigate the mobile genetic elements underlying chromosomal *bla*_CTX-M_ integration, we characterized their associated insertion structures. (**Supplementary Note 3**) Among the 864 chromosomal *bla*_CTX-M_, 668 (77.3%) were associated with IS*Ecp1*, 163 (18.9%) with IS*26* and 33 (3.8%) with other mobile genetic elements or unresolved architectures. Among the four predominant variants: 70.2% and 26.2% of strains carrying *bla*_CTX-M-15_ were associated with ISEcp1 and IS26 integrations, respectively, versus 91.6% and 4.9% for *bla*_CTX-M-14_, 86.5% and 9.5% for *bla*_CTX-M-55_ and 66.7 and 22.2% for *bla*_CTX-M-27_. Multiple chromosomal copies of the same *bla*_CTX-M_ variant were predominantly IS*Ecp1*-associated (64/72; 89%) (**Extended Data Table 1; Extended Data Fig. 7; Supplementary Fig. S4**). The relative IS contribution varied across STs: IS*Ecp1*-associated integrations predominated in ST38 (100%; 102/102), ST405 (100%; 40/40), ST10 (91.9%; 34/37) but was less prevalent in ST131 (65.8%; 146/222). Within ST131, IS*Ecp1* predominated in clades A (100%, 46/46) and C1 (87.5%, 35/40), whereas IS26-associated structures predominated in C2 (56.2%, 68/121).

IS*26*-associated insertions carried significantly more ARGs than IS*Ecp1*-associated ones (mean 2.48 *vs.* 0.85 additional resistance genes; difference 1.64 [95% CI 1.33–1.94], p<0.001). Among the two most frequent barcodes, [*aac*(3)-IIa]_[*bla*_OXA-1_]_[*aac*(6’)-Ib-cr] was predominantly IS*26*-mediated (81.3%, 61/75), while [*qnrS1*] alone was almost exclusively IS*Ecp1*-mediated (91.2%, 31/34) (**Fig.3d-g; Extended Data Fig. 7**).

IS*Ecp1* was the dominant driver of chromosomal *bla*_CTX-M_ integration across variants and STs, mediating compact insertion events, while IS*26*, though less frequent, assembled larger and more complex multidrug-resistance structures.

### Chromosomal *bla*_CTX-M_ integration is broadly distributed across the *E. coli* chromosome, with ST-dependent integration sites

Using a synteny-based computational approach, chromosomal insertion sites were successfully mapped onto the *E. coli* K-12 reference genome for 95.9% (829/864) of *bla*_CTX-M_ loci. Dual-flank resolution was high for *bla*_CTX-M-15_ and *bla*_CTX-M-27_ (79.5% and 83.8%, respectively) consistent with simple, structurally well-defined insertion events, whereas it was lower for *bla*_CTX-M-55_ (49.4%). This lower resolution was not explained by assembly quality but was associated with frequent discrepancies between upstream and downstream coordinates (45.3% vs. 9.5% for *bla*_CTX-M-15_), suggesting more complex chromosomal rearrangements around *bla*_CTX-M-55_.

Application of this mapping strategy revealed that chromosomal insertions were distributed across the entire *E. coli* K-12 backbone, with no protected zones or large genomic regions refractory to integration. Despite this broad chromosomal permissiveness, insertion events were not uniformly distributed: distinct hotspots emerged, strongly associated with specific genetic background/ *bla*_CTX-M_ variant combinations. Most notably, *bla*_CTX-M-15_ in ST131 clustered around 48 and 52 min, *bla*_CTX-M-27_ in ST131 around 75 min, and bla_CTX-M-14_ in ST38 around 55 min and 78–83 min. These positional clusters likely reflect either recurrent independent integration at preferred chromosomal targets or clonal expansion following a single early integration event **(Fig.4).**

**Figure 4.**
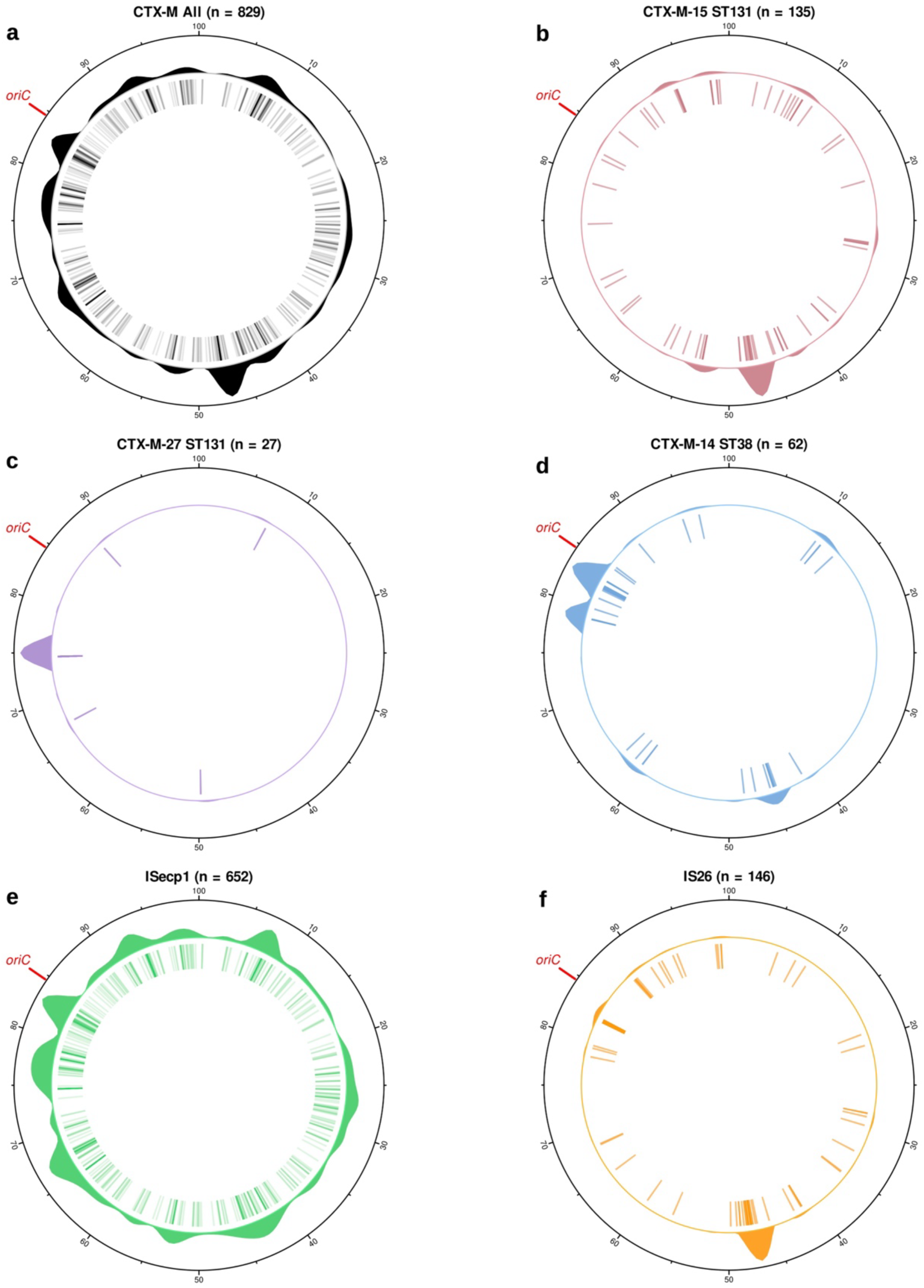
Chromosomal distribution of *bla*_CTX-M_ integration sites in *Escherichia coli*. Circular plots show the genomic distribution and recurrence of chromosomal *bla*_CTX-M_ integration sites. The outermost circle represents the *E. coli* chromosome on a 0–100 min scale. The inner circle indicates the chromosomal position of individual integration events, whereas the outer histogram represents the number of genomes with a *bla*_CTX-M_ insertion at each position, highlighting recurrent integration sites. a Distribution of all chromosomal *bla*_CTX-M_ insertions identified in the dataset (n = 829). b, Distribution of *bla*_CTX-M-15_ integration sites within ST131 (n = 135). c, Distribution of *bla*_CTX-M-27_ integration sites within ST131 (n = 27). d, Distribution of *bla*_CTX-M-14_ integration sites within ST38 (n = 62). e-f, distribution of *bla*_CTX-M_ integration sites depending on IS type integration ISEcp1 (n = 652) or IS26 (n=146).

## DISCUSSION

Chromosomal integration of *bla*_CTX-M_ *in ESBL*-*E. coli* emerges from this large-scale analysis as a frequent and lineage-structured process, rather than an anecdotal endpoint of ESBL evolution. The spread of CTX-M enzymes has largely been interpreted as a plasmid-driven epidemic^4,8^, in which conjugative plasmids enabled rapid horizontal dissemination across lineages, hosts and reservoirs. Our findings support a complementary model in which plasmids remain central to early *bla*_CTX-M_ dissemination and chromosomal integration represents a secondary evolutionary route through which resistance modules become stabilized within successful *E. coli* backgrounds. Once embedded in the chromosome, *bla*_CTX-M_ and co-integrated ARGs could become less horizontally mobile but be maintained through vertical inheritance within successful lineages.

Chromosomal *bla*_CTX-M_ has been reported in human, animal or environmental *E. coli* collections across Europe, Japan, and Africa, with recurrent involvement of ST38 and ST131 ^17–19,22,23,30,31^. Our analysis extends these observations showing that chromosomal integration is a widespread and structured component of the CTX-M epidemic.

ST38 has emerged as the clearest example of lineage-associated chromosomal integration displaying the highest integration rate among major STs and a strong association with chromosomal *bla* ^32^ This is consistent with reports of chromosomally encoded ESBL genes in ST38 clinical and wildlife isolates. ST38 has also been implicated in chromosomal integration of *bla*_OXA-244_ and *bla*_OXA-48-like_ structures^33,34^. A recent study showed that the antiplasmid system ApsAB promoted *bla*_OXA-48_ chromosomal integration and plasmid elimination in ST38^33^, suggesting that this background may be particularly permissive to chromosomal integration of acquired resistance determinants.

ST131 shows a distinct pattern combining substantial chromosomal integration with unparalleled global dissemination. In our dataset, as expected, it was the dominant lineage^8,35^ and accounted for a large share of chromosomal *bla*_CTX-M-15_ and *bla*_CTX-M-27_ integrations. Previous studies have reported contrasting subclade-specific trajectories. In Japan, *bla*_CTX-M-14_ dissemination within C1-ST131 shifted from conjugative plasmid spread to clonal expansion following a single ISEcp1-mediated chromosomal integration event^36^, while in Hungary, *bla*_CTX-M-27_ dissemination in C1-ST131 remained plasmid borne, whereas *bla*_CTX-M-15_ in C2 was predominantly chromosomally integrated^37^. Our broader analysis confirms marked subclade heterogeneity, with integration rates ranging from 20% in C1 to 60% in C2, despite their relatively recent divergence^38^. Exploratory modelling, was consistent with a more recent increase in C2, suggesting that chromosomal integration may have contributed to vertical inheritance of *bla*_CTX-M-15_ during C2 clonal expansion.

This lineage-dependent structuring of chromosomal *bla*_CTX-M_ integration echoes observations made of virulence-associated genes in *E. coli*, where the genomic location of iron acquisition systems was shown to depend on strain phylogeny^39^, suggesting strong epistatic interactions shaping adaptive gene fixation at the genomic scale.

Chromosomal *bla*_CTX-M_ are co-located with additional resistance genes in approximately one third of genomes. The diversity and limited sharing of ARG barcodes between CTX-M variants support multiple independent integration events from distinct genetic structures. These co-located genes confer resistance to several antibiotic classes, including aminoglycosides and fluoroquinolones, potentially favoring co-selection and persistence of ESBL-*E. coli* ^30,40^.

Two mobile-element architectures dominated chromosomal *bla*_CTX-M_ integration. ISEcp1 can mobilize *bla*_CTX-M_ together with variable amounts of adjacent DNA^41^ and was originally involved in mobilizing *bla*_CTX-M_ genes from the chromosome of environmental *Kluyvera* spp. onto mobile elements^12,42^. IS26 promotes integration, amplification and rearrangement of larger ARG-rich regions^43,44^. Accordingly, multicopy *bla*_CTX-M_ loci may result from repeated ISEcp1-mediated insertion events or IS26-associated amplification^19,45,32^ . Together, these mechanisms provide a link between horizontal acquisition and subsequent chromosomal integration of ESBL resistance determinants.

While previous studies on limited collections from Niger^23^ and Madagascar^22^ suggested that chromosomal *bla*_CTX-M_ insertions spared the *Ori* macrodomain, our large-scale dataset nuances this picture: integration occurs across the entire *E. coli* chromosome without evidence of large regions refractory to integration. However, integration was not random, with distinct ST/variant-specific integration sites suggesting either that the genetic environment may have favored the integration at this exact location or that the insertion was followed by a successful clonal expansion.

The evolutionary consequence of chromosomal integration is uncertain. Plasmid-borne *bla*_CTX-M_ is usually mobile and highly transferable but plasmids may be costly, unstable or subject to host restriction ^8,16^. Chromosomal *bla*_CTX-M_ is less horizontally mobile, once integrated into a successful clone, it may be maintained through vertical inheritance without requiring plasmid persistence. These dissemination modes are not mutually exclusive. Chromosomal integration can promote vertical inheritance of *bla*_CTX-M_ within successful lineages, while these backgrounds continue to acquire and exchange plasmid-borne resistance determinants. Conversely, chromosomal *bla*_CTX-M_ regions may be remobilized onto plasmids together with associated ARGs^14,43,44^. Chromosomal integration should therefore not be interpreted as the end of *bla*_CTX-M_ mobility, but as a shift in its epidemiological risk profile, combining clonal persistence with continued opportunities for horizontal resistance acquisition.

Several limitations should be acknowledged. Public long-read or hybrid assemblies are subject to sampling or deposition biases limiting estimates of global prevalence. Nevertheless, the large non-redundant dataset spanning 666 BioProjects provides a broad picture of chromosomal *bla*_CTX-M_ integration across the global *E. coli* population. Limited source, geographic, and temporal metadata restricted epidemiological analyses, and the ST131 subclade integration-rate model should thus be considered exploratory rather than predictive.

Future studies should experimentally address the stability, expression, fitness cost, and transferability of chromosomal *bla*_CTX-M_ across distinct genomic contexts and identify conditions favouring chromosomal integration. Longitudinal studies will also be essential to determine the role that chromosomal integration might play in the future trajectory of ESBL-*E. coli* and antimicrobial resistance more broadly.

Our findings have direct implications for AMR surveillance. Beyond resistance genes and sequence types, genomic location, insertion sites, flanking IS elements and co-integrated ARG barcodes should also be monitored, as these features can help resolve dissemination routes, identify co-selected resistance determinants, and better understand AMR dynamics. Once established in successful lineages such as ST38 or ST131, chromosomal *bla*_CTX-M_ disseminates efficiently through clonal expansion ^6–8^, demonstrating that reduced horizontal mobility does not imply reduced epidemiological impact.

More broadly, our large-scale genomic analysis raises a fundamental evolutionary question: are ESBL-*E. coli* progressively domesticating *bla*_CTX-M_ as a stable chromosomal trait, echoing *Kluyvera* spp., where CTX-M-like enzymes (KLU) are natively chromosomal? If so, this would mark a shift from a plasmid-driven epidemic toward one shaped by chromosomal entrenchment, making ESBL resistance increasingly embedded within *E. coli* population structure and less dependent on plasmid transmission **(Fig. 5).** Whether carbapenemase genes will follow the same evolutionary path remains a critical question for anticipating the next phase of the AMR crisis.

**Figure 5.**
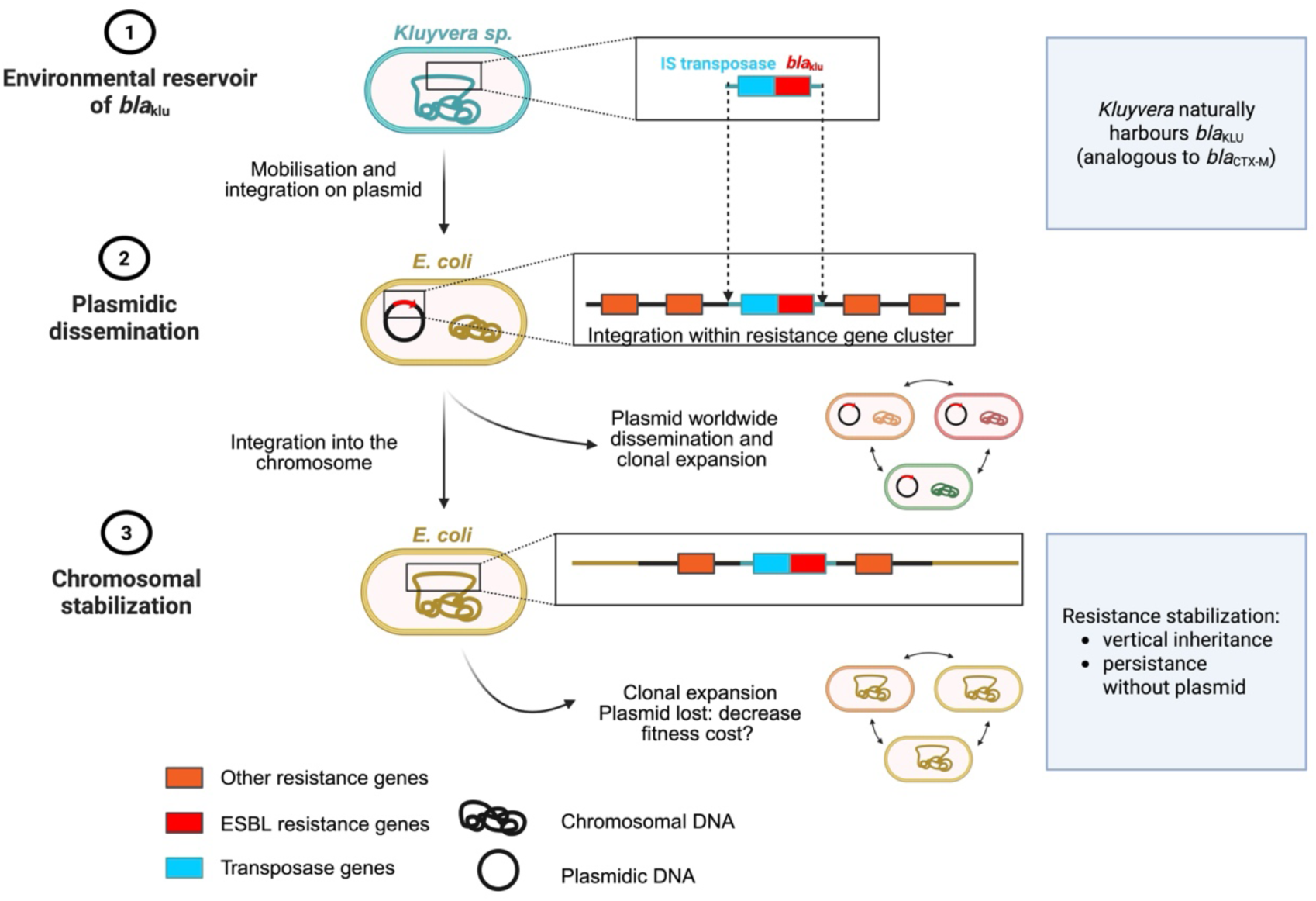
Proposed natural history of *bla*_CTX-M_ mobilization and chromosomal integration. Schematic representation of the proposed evolutionary pathway of *bla*_CTX-M_ genes, originating from probable ancestral chromosomal KLU-type β-lactamases in *Kluyvera* spp., followed by their mobilization by insertion sequences, transfer onto mobile genetic elements, evolutionary diversification and subsequent integration into the chromosome of *Escherichia coli*. The model also illustrates the progressive acquisition and co-mobilization of additional antimicrobial resistance determinants during this process. *bla*_CTX-M_ genes are shown in red, other antimicrobial resistance genes in orange and transposase genes in blue. Created in BioRender. Agreement number SE2A9WS1M0, Berti, V. (2026) https://BioRender.com/d9y3uqj.

## MATERIALS AND METHODS

### Genome collection and inclusion criteria

In March 2026, we queried the NCBI Genomes database to identify all available *Escherichia coli* genomes. Because accurate assignment of *bla*_CTX-M_ genes to plasmid or chromosomal locations requires resolved genomic context, we restricted the analysis to genomes generated using long-read or hybrid-assembly approaches for which both fasta and GBFF files were available. Based on NCBI data, genomes were retained if they met the following criteria: taxonomic identifier 562 (*E. coli*), Average Nucleotide Identity check status reported as “OK”, assembly N50 ≥100 kb, genome completeness ≥90%, contamination ≤5%, and an available BioProject accession. Genomes lacking a BioProject accession or release date were excluded.

*Bla*_CTX-M_ genes were detected using ResFinder, with minimum identity and coverage thresholds of 80%. Genomes carrying at least one *bla*_CTX-M_ gene were retained as CTX-M-producing ESBL-*E. coli*. To minimize overrepresentation of outbreak-associated or repeatedly deposited genomes, closely related duplicates were removed. Genomes were considered duplicates when all three of the following criteria were met: (i) they belonged to the same BioProject; (ii) their core-genome MultiLocus Sequence Typing (cgMLST) distance was ≤10 allelic differences ^46^; and (iii) they had identical antimicrobial resistance gene profiles. Core-genome MLST was performed using chewBBACA v3.5.2 through the AlleleCall pipeline, based on the *E. coli* stable cgMLST Achtman scheme comprising 2,513 core genes, and pairwise allelic distances were computed using cgmlst-dists v0.4.0 (https://github.com/tseemann/cgmlst-dists). After duplicate removal, a single representative genome was retained per cluster, yielding the final non-redundant genome collection used for downstream analyses.

Genome metadata were collected from NCBI, and EnteroBase when available. The full genome selection workflow is detailed in **Extended Data Fig. 1**. All genome data and metadata generated or analyzed in this study are available in the Zenodo repository (https://doi.org/10.5281/zenodo.20934804).

### Genome typing and antimicrobial resistance gene detection

Phylogroups were assigned using ClermontTyper. Sequence types (STs) were determined using MLST according to the Achtman scheme implemented in MLST v2.23 (https://github.com/tseemann/mlst). O-serotypes and fimH alleles were identified using ABRicate v1.0.1 (https://github.com/tseemann/abricate) with the EcOH and FimTyper databases, respectively. Antimicrobial resistance genes, including *bla*_CTX-M_ variants and carbapenemase genes, were detected using ABRicate v1.0.1 with the ResFinder database. Insertion sequences were detected using ABRicate v1.0.1 with the ISFinder database (October 2020 version).

For ST131 genomes, subclades A, B, C1 and C2 were assigned based on O-type, *fimH*-type, *bla*_CTX-M_ variant and previously described ST131 subclade definitions ^47^. Clonal complexes were retrieved from MLST or EnteroBase classification where available.

### Assignment of *bla*_CTX-M_ genomic location

The plasmid or chromosomal origin of each contig was inferred using contigs size and Platon v1.7 ^26^. Each detected *bla*_CTX-M_ gene was assigned to a plasmid-borne or chromosomal location based on the classification of its host contig. As large contigs may occasionally be misclassified, contigs longer than 500 kb were manually reviewed. Overall, 94.1% (2,295/2,440) of chromosomal contigs were above the 500-kb threshold, including 94.3% (725/769) of *bla*_CTX-M_-carrying chromosomal contigs. All plasmidic contigs were below 500 kb.

Analyses were performed at two levels. At the gene level, each *bla*_CTX-M_ copy was considered independently, allowing genomes carrying multiple copies to contribute more than one event. At the genome level, a genome was classified as carrying chromosomal *bla*_CTX-M_ when at least one copy was assigned to the chromosome. Genomes carrying both chromosomal and plasmid-borne *bla*_CTX-M_ copies were retained in the dataset and analysed according to the location of each individual *bla*_CTX-M_ gene.

### Descriptive analyses of chromosomal *bla*_CTX-M_ integration

The distribution of *bla*_CTX-M_ genes was described by genomic location, phylogroup, ST, clonal complex and CTX-M variant. For each category, the chromosomal integration rate was calculated as the proportion of *bla*_CTX-M_ genes assigned to the chromosome relative to the total number of *bla*_CTX-M_ genes detected in that category, with confidence intervals estimated for each proportion. Genome-level integration rates were calculated as the proportion of genomes carrying at least one chromosomal *bla*_CTX-M_ gene within each subgroup.

To assess whether chromosomal integration patterns were driven by clonal overrepresentation or specific resistance backgrounds, sensitivity analyses were performed. First, chromosomal integration rates were compared between the non-redundant dataset and the set of related or duplicate genomes identified before deduplication. Second, ST-level analyses were repeated after exclusion of carbapenemase-producing genomes, defined as genomes carrying at least one acquired carbapenemase gene (**Supplementary Fig. S3**). Third, within high-risk clones, chromosomal integration rates were compared between ESBL-only and carbapenemase-harboring genomes using Fisher’s exact test (significance threshold p<0.05). These analyses were used to determine whether the distribution of chromosomal *bla*_CTX-M_ integration across major STs persisted after accounting for related genomes and after removing carbapenemase-harboring genomes (**Supplementary Fig. S3)**.

### Pangenome analysis and phylogenetic reconstruction

Pangenome analysis was performed using PPanGGOLiN v2.2.4 to define the persistent genome partition of the *E. coli* population. Maximum-likelihood phylogenetic inference was conducted from the persistent genome alignment using FastTree v2.10.0 under a generalized time-reversible model with gamma correction. The phylogeny was rooted using an *Escherichia fergusonii* outgroup (GCA_020097475.1 ^35^) and visualized using iTOL. Branches were colored according to phylogroup. Metadata rings indicated, from inner to outer ring, chromosomal *bla*_CTX-M_ variants, plasmid-borne *bla*_CTX-M_ variants, and the ten most prevalent STs.

### Statistical analyses for lineages chromosomal integrations

Statistical analyses were performed using R. Chromosomal integration was analysed as a binary outcome at the *bla*_CTX-M_ gene level, with chromosomally located genes coded as 1 and plasmid-borne genes coded as 0. Logistic regression models with a binomial error structure were used to estimate average chromosomal integration rates according to sequence type, CTX-M variant, CTX-M family, and ST131 subclade.

For ST-level analyses, only the most prevalent sequence types were included in formal comparisons. Estimated chromosomal integration probabilities and 95% confidence intervals were obtained on the response scale. Expected numbers of chromosomal *bla*_CTX-M_ genes for each group were calculated by multiplying the estimated chromosomal integration probability by the total number of *bla*_CTX-M_ genes observed in that group. Pairwise comparisons between sequence types were performed using Tukey-adjusted post-hoc tests, and differences were summarized using compact letter displays. Model estimates, standard errors and p-values are reported in **Supplementary Table S2**.

Within ST131, chromosomal integration rates were compared across subclades A, B, C1 and C2 using logistic regression, with C1 as the reference category. Estimates, standard errors and p-values are reported in **Supplementary Table S3**.

### Modelling chromosomal *bla*_CTX-M_ accumulation within ST131

Chromosomal *bla*_CTX-M_ accumulation within ST131 was approximated using an exploratory model based on the dated phylogenetic structure of subclades A, B, C1 and C2. The model assumed a linear increase over time in the logit-transformed integration rates of *bla*_CTX-M_ genes, with branch-specific accumulation speeds estimated across the A/B, B/C and C1/C2 divergences. Current subclade-specific integration rates were derived from logistic regression estimates, and uncertainty in both integration rates and divergence dates was propagated through one million simulations. Projected trajectories from 1980 to 2025 were generated to assess whether present-day integration patterns were compatible with changes in accumulation speed during ST131 diversification. The full model is described in **Supplementary Note 1 and Extended Data Fig. 4.**

### Multicopy and multi-integration analyses

Genomes carrying more than one chromosomally assigned *bla*_CTX-M_ gene were analysed separately. Multicopy integration was defined as the presence of more than one chromosomal copy of the same *bla*_CTX-M_ variant in a single genome and multi-integration as the presence of chromosomal copies of two or more distinct *bla*_CTX-M_ variants in the same genome. For these genomes, *bla*_CTX-M_ variants, sequence types and ARG barcodes were described to determine whether multicopy and multi-integration events were restricted to specific lineages or occurred across multiple genetic backgrounds.

### Definition of variant-specific genomic windows around chromosomal *bla*_CTX-M_ genes

To characterize the genomic environment surrounding chromosomally integrated *bla*_CTX-M_ genes, we defined variant-specific windows for the four most frequent chromosomal *bla*_CTX-M_ variants (*bla*_CTX-M-15_, *bla*_CTX-M-55_, *bla*_CTX-M-14_ and *bla*_CTX-M-27_) capturing the relevant inserted sequence while avoiding arbitrary flanking distances (See **Supplementary Note 2**)

For each variant, sequences located up to 100 kb upstream and 100 kb downstream of *bla*_CTX-M_ were extracted from genomes carrying the corresponding variant and were analyzed separately. Mash-based phylogenetic trees were generated using Mashtree v1.4.6, and patristic distances were computed for every genome pair carrying the same *bla*_CTX-M_ variant. For each variant and direction, k-means clustering was applied to the distribution of patristic distances. The number of clusters was selected as the smallest value for which the ratio of between-cluster to total sum of squares exceeded 95% (**Supplementary Table S4**). The first cluster, corresponding to the shortest patristic distances, was interpreted as containing related genomic contexts. The upper boundary of this cluster was used as the threshold above which pairs of genomic contexts were considered unrelated (**Supplementary Figure S2**).

Using these variant- and direction-specific thresholds, patristic distances between all pairs of unrelated genomes were recalculated using window sizes ranging from 1 to 100 kb, increasing in 1-kb increments from the *bla*_CTX-M_ gene. For each genome pair, the minimum window size was defined as the shortest distance from *bla*_CTX-M_ at which the patristic distance exceeded the unrelated-genome threshold. The final window retained for each variant corresponded to the smallest upstream/downstream interval for which at least 95% of unrelated genome pairs had a patristic distance above the threshold. The resulting windows were −16 to +33 kb for *bla*_CTX-M-15_, −24 to +5 kb for *bla*_CTX-M-55_, −9 to +14 kb for *bla*_CTX-M-14_, and −15 to +21 kb for *bla*_CTX-M-27_, where negative and positive values indicate positions upstream and downstream of *bla*_CTX-M_, respectively. The detailed procedure and thresholds are provided in **Supplementary Note 2, Supplementary Table S4, and Supplementary Fig. S2.**

### ARG barcode definition and genetic-environment analysis

Variant-specific windows were used to characterize the antibiotic resistance-gene (ARG) content surrounding chromosomally integrated *bla*_CTX-M_ genes. Within each window, ARGs were identified using ResFinder. For each *bla*_CTX-M_ insertion, the ordered combination of non-*bla*_CTX-M_ ARGs located upstream and downstream of *bla*_CTX-M_ was recorded in accordance with the Open Reading Frame of the *bla*_CTX-M_ gene. This ordered combination was defined as an ARG barcode.

ARG barcodes were compared within and between *bla*_CTX-M_ variants to determine whether chromosomal integrations shared conserved resistance architectures or represented variant-specific genetic environments. Barcodes were summarized by *bla*_CTX-M_ variant, sequence type and recurrence across genomes. Shared, plasmid-borne co-localized and variant-specific barcodes were used to assess whether chromosomal *bla*_CTX-M_ integration corresponded to isolated *bla*_CTX-M_ insertion or to stabilization of broader multidrug-resistance islands.

Chromosomal barcodes identical to a plasmidic barcode, or contained within a larger plasmidic barcode, were used to assess whether chromosomal *bla*_CTX-M_ integration corresponded to isolated gene insertion or to stabilization of broader multidrug-resistance islands.

The complete ARG barcode table is provided in **Supplementary Table S2-c**.

### Representative ISEcp1- and IS26-associated chromosomal integration structures

To explore the mechanisms underlying chromosomal *bla*_CTX-M_ integration, representative insertion structures were selected from recurrent chromosomal integration patterns, focusing on two major mobile-element architectures: ISEcp1, predominantly associated with *bla*_CTX-M_ alone, and IS26, associated with ARG-rich composite structures. For each representative structure, the local genetic organization was reconstructed from assembly and annotation files, including *bla*_CTX-M_, flanking ARGs, insertion sequences, mobile element orientation, flanking chromosomal genes, and insertion boundaries. For each *bla*_CTX-M_ variant, genomic windows flanking the gene were extracted and aligned by synteny analysis using Clinker to identify conserved sequence motifs. Insertion sequences shared across multiple strains were classified into three categories according to the suspected transposition mechanism described in **Supplementary Note 3**. A subset of the most frequent insertion events, was manually re-annotated using SnapGene, ISfinder and TnCentral softwares to precisely delineate insertion boundaries. Together, these analyses were used to determine whether recurrent chromosomal *bla*_CTX-M_ integrations were compatible with distinct mechanistic routes of chromosomal stabilization.

### Mapping of chromosomal *bla*_CTX-M_ insertion sites

For chromosomally assigned *bla*_CTX-M_ genes, insertion contexts were characterized from annotated GBFF files. The closest upstream and downstream *E. coli* chromosomal genes flanking the *bla*_CTX-M_-containing insertion were identified, excluding genes belonging to the inserted region where possible. These flanking genes were used as genomic anchors to localize insertion sites within the *E. coli* chromosome.

For each locus, up to five upstream and five downstream CDS features were mapped to the *E. coli* K-12 reference chromosome ^48^ using gene symbols, locus tags, and product descriptions. Coordinates were expressed on the conventional 0–100 min chromosomal scale, with circular distances calculated using modulo 100 arithmetic. To minimize the impact of chromosomal rearrangements and misannotations, a dynamic programming algorithm identified the longest contiguous chain of anchor genes on each flank (maximum gap ⩽ 1.0 min; minimum two genes required).

Insertion coordinates (rounded to 0.1 min) were assigned based on the available flanking information. When valid chains were identified on both flanks and the junction gap between the closest anchors was ≤5.0 min, the insertion coordinate was defined as their circular midpoint. When the junction gap exceeded 5.0 min, the flank with the longer valid chain was retained; when only one flank was valid, the position of its nearest anchor was used. Loci lacking a stable anchor chain, or with equally supported but discordant flanks, were classified as not determinable (ND).

Mapped insertion sites were visualized using R on circular representations of the *E. coli* chromosome, with individual integration positions shown on the inner circle and the number of genomes carrying an insertion at each position, represented by an outer histogram. Genomes sharing the same flanking-gene pair were considered to belong to the same integration region. Distributions were examined globally and for selected variant–lineage combinations (ST131-*bla*_CTX-M-15_, ST131-*bla*_CTX-M-27_ and ST38-*bla*_CTX-M-14_) to identify recurrent insertion regions and assess their association with specific *bla*_CTX-M_ variants or *E. coli* lineages.

All softwares and databases are listed in the Supplementary Information.

## Supporting information

Supplementary References

## Extended Data

**Extended Data Figure 1.**
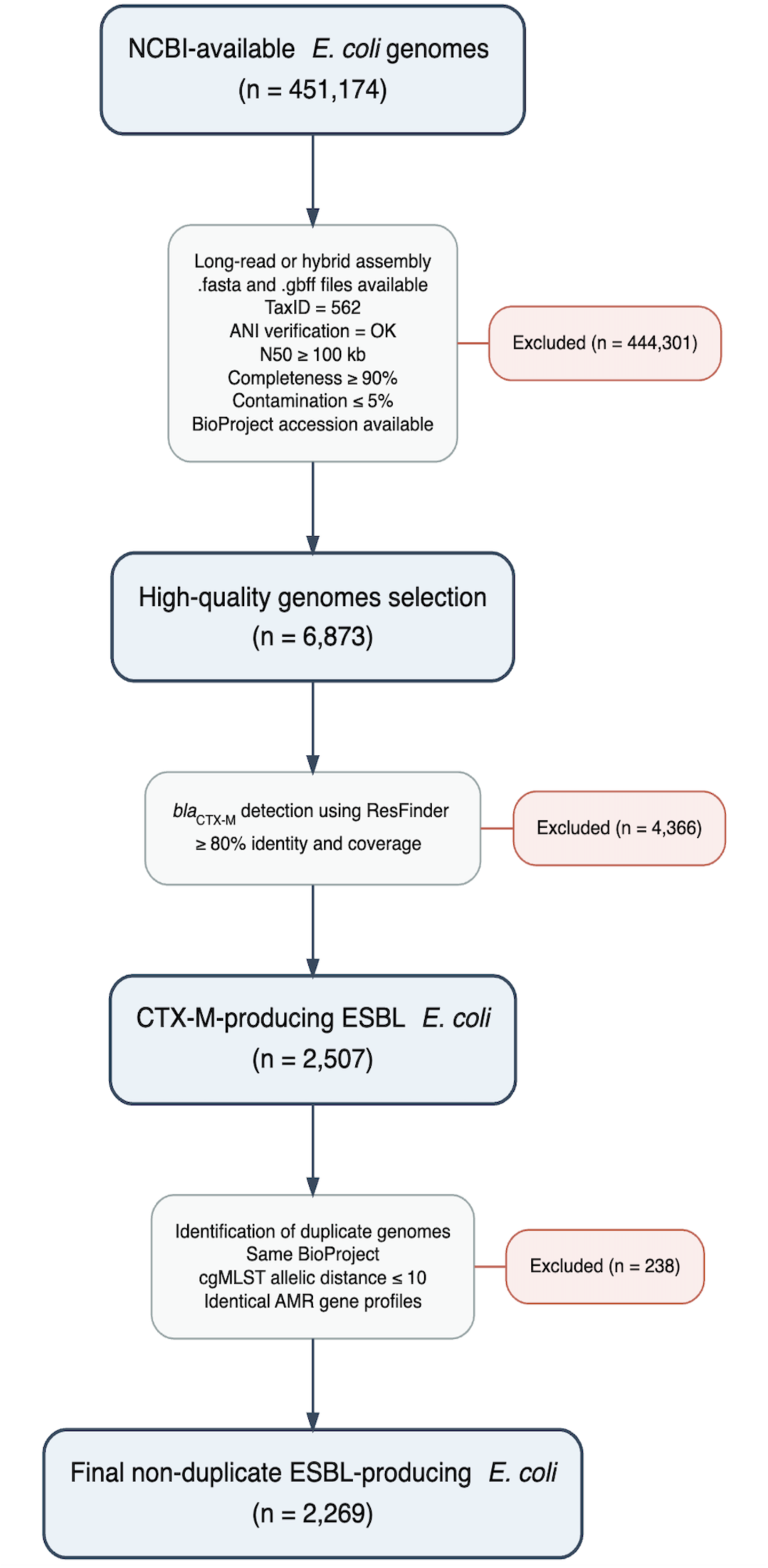
Selection workflow of the ESBL-producing *E. coli* genome collection included in this study. Starting from 451,174 NCBI-available genomes, quality filtering retained 6,873 high-quality genomes. Screening for *bla*_CTX-M_ genes using ResFinder database identified 2,507 genomes. After removal of duplicate genomes based on BioProject origin, cgMLST similarity (≤10 allelic differences).

**Extended Data Fig. 2.**
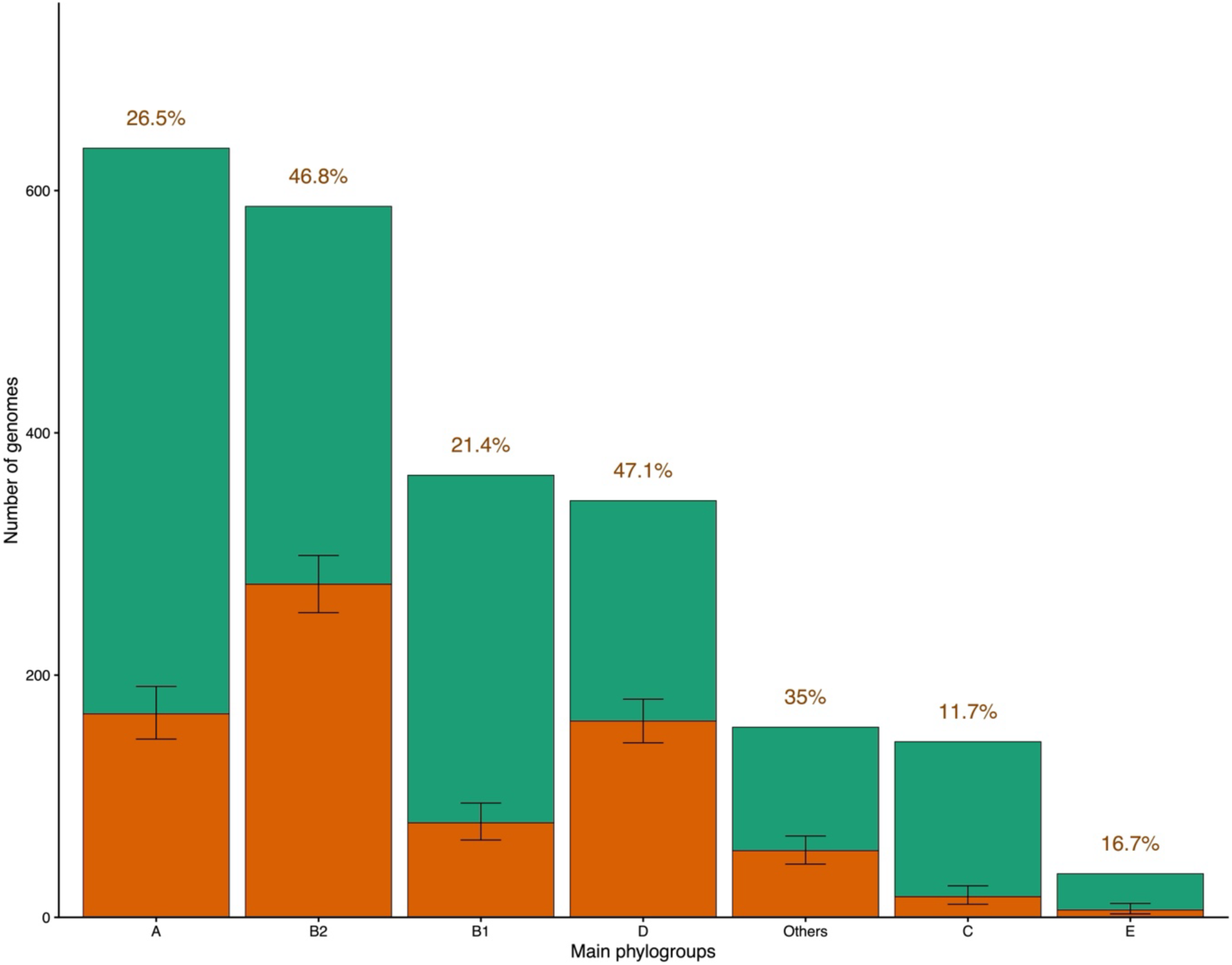
Distribution of chromosomal *bla*_CTX-M_ gene integration among the main *E. coli* phylogroups. Bars represent the number of genomes carrying at least one chromosomal *bla*_CTX-M_ gene (orange) or no chromosomal *bla*_CTX-M_ gene (green). Total bar height corresponds to the number of *bla*_CTX-M_-carrying genomes identified in each phylogroup. Percentages displayed above each bar indicate the proportion of genomes carrying at least one chromosomal *bla*_CTX-M_ gene within the corresponding phylogroup. Error bars represent the 95% confidence interval of this proportion. Chromosomal and plasmid locations were inferred *in silico* from genome assembly analyses.

**Extended Data Figure 3.**
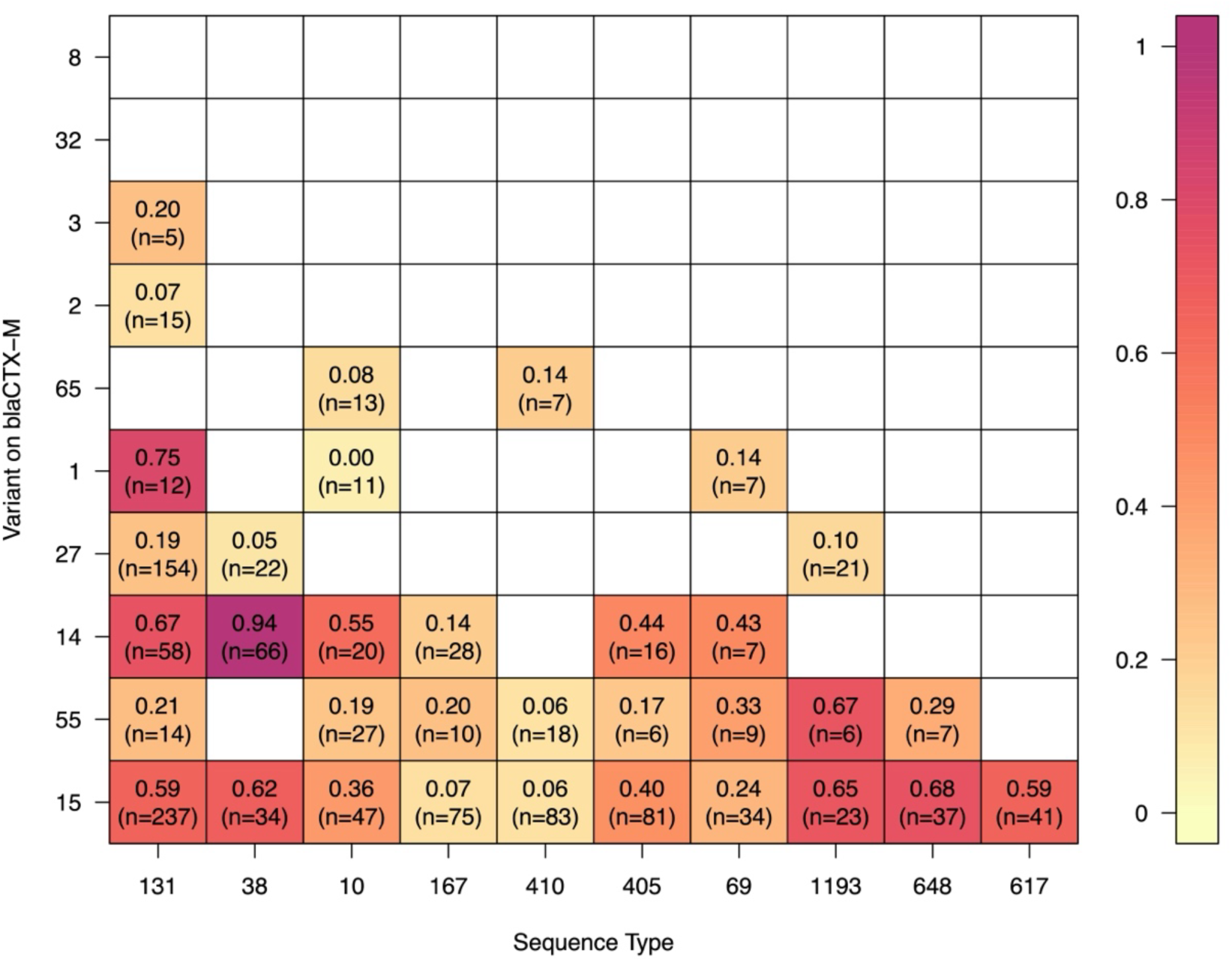
Heatmap of the integration rates for couples of a given Sequence Type and given variant of *bla*_CTX-M_. A couple is displayed if it is observed at least five times in the dataset. The integration rate and the number of occurrences of the couple are indicated for each displayed couple. Only the ten main ST and the ten main variants are displayed.

**Extended Data Figure 4.**
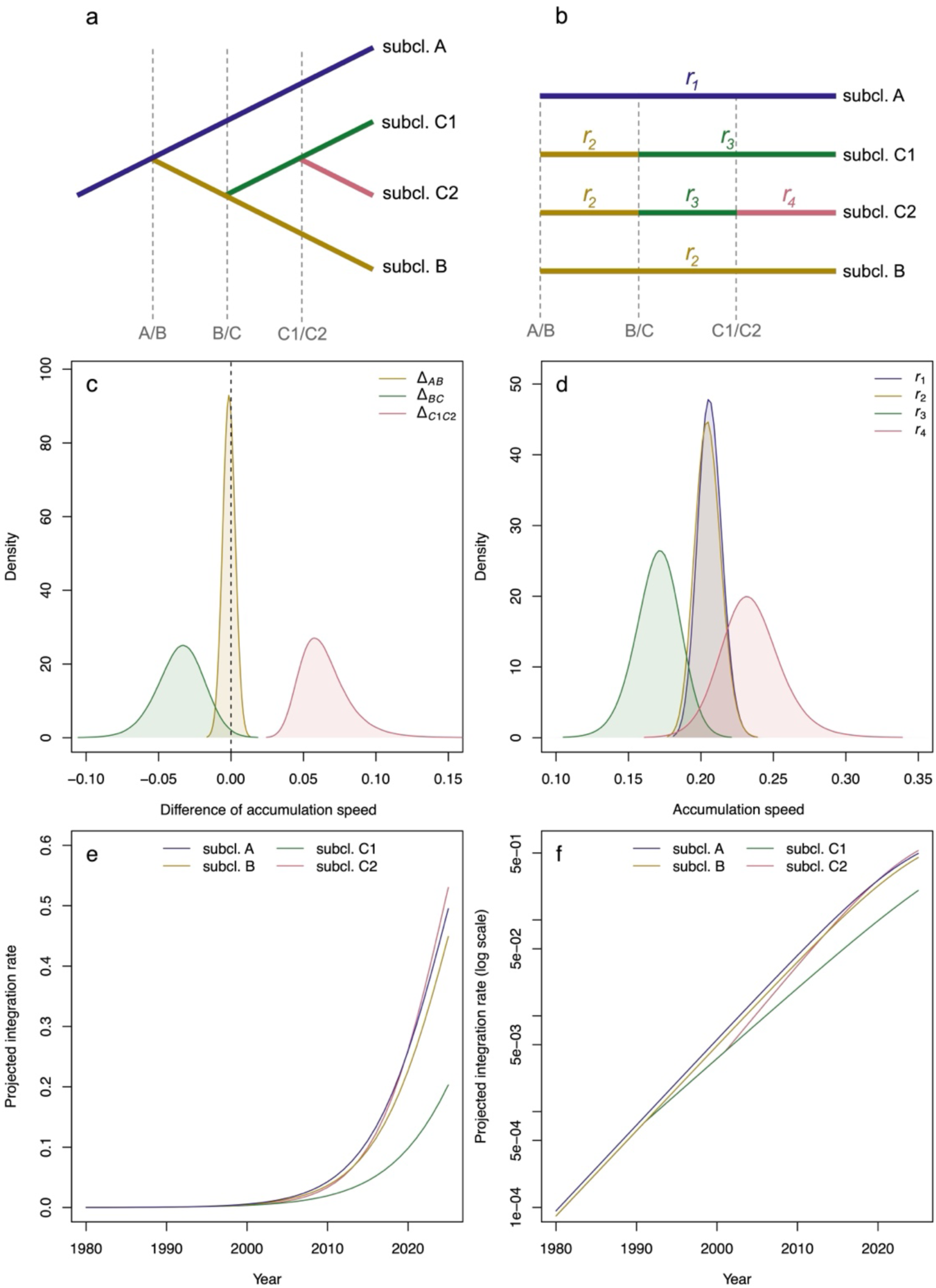
Model of integration accumulation speed in ST131. In **a,** schematic representations of the phylogenetic structure of the four ST131 subclades A, B, C1 and C2. **b,** Variations in accumulation speed in the subclades over time. **a-b**, The branch leading to subclade A (purple) is associated with *r*_1_ , the branch diverging at A/B and leading to subclade B (yellow) with *r*_2_, the branch diverging at B/C and leading to subclade C1 (green) with *r*_3_, and the branch diverging at C1/C2 and leading to subclade C2 (pink) with *r*_4_. **c,** Distribution of the changes in accumulation speed at the A/B divergence (Δ*_AB_*, yellow), at the B/C divergence (Δ*_BC_*, green), and at the C1/C2 divergence (Δ*_C1C2_*, pink). **d,** Distribution of accumulation speeds projected by the model, accounting for uncertainty on current integration rates and divergence dates. **e-f,** Projected variation in the chromosomal integration of *bla*_CTX-M_ according to the model from 1980 to 2025, on a linear (**e**) and log-scale (**f**). **d-f,** Values estimated from the model using *Y_A_*(*t_AB_*) = 10^-12^ (Supplementary note 1).

**Extended Data Figure 5.**
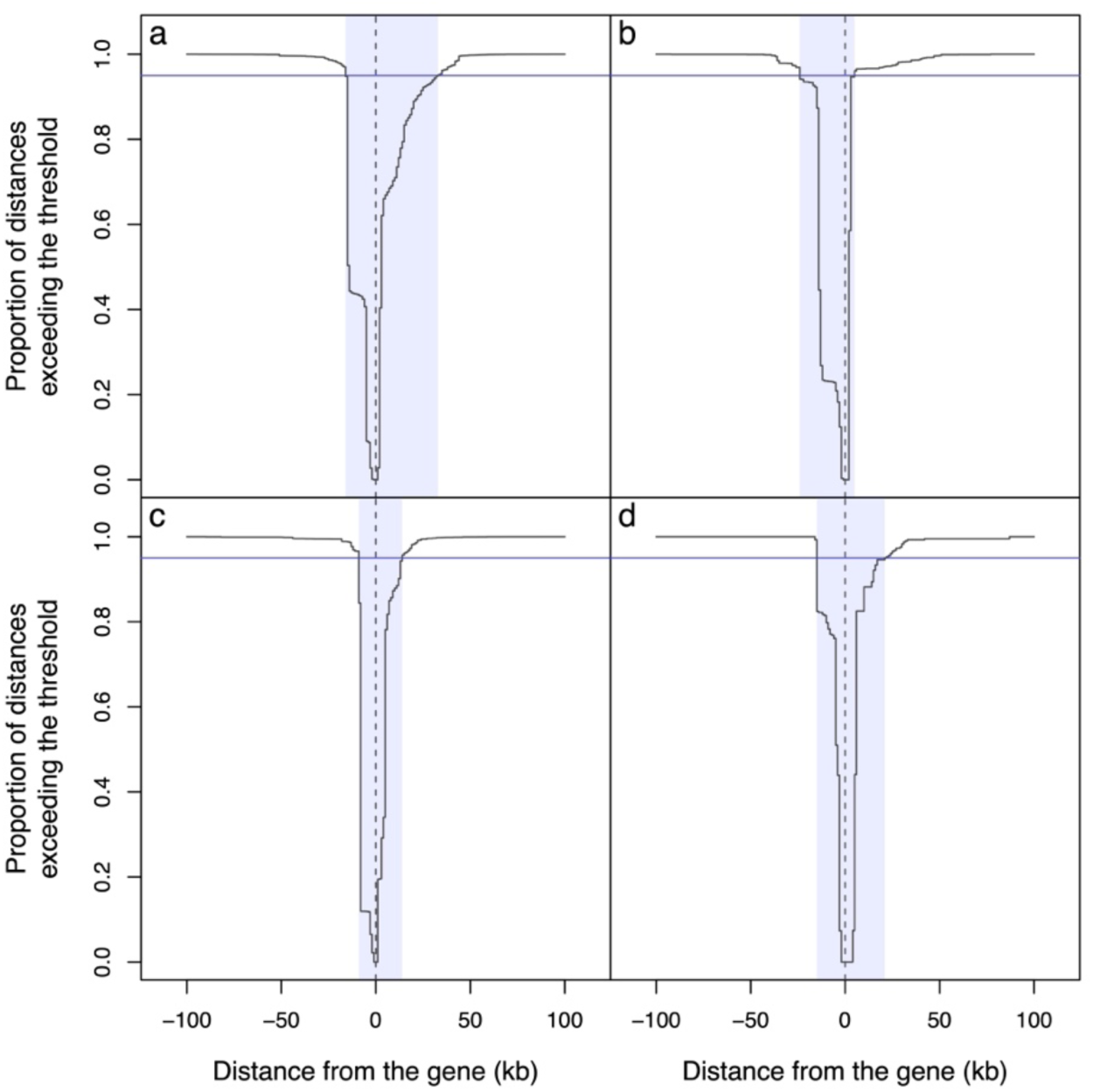
Delineation of the variant-specific minimal windows. Proportion of distances greater than the threshold between pairs of unrelated genomes carrying the variant CTX-M-15 (**a**), CTX-M-55 (**b**), CTX-M-14 (**c**), CTX-M-27 (**d**). The zero of the X-axis corresponds to the position of the *bla*_CTX-M_ gene (dashed vertical line). Negative distances are upstream of the gene and positive distances are downstream of the gene. The window for each panel (blue shade) corresponds to 95% of distances above the threshold (blue line).

**Extended Data Figure 6.**
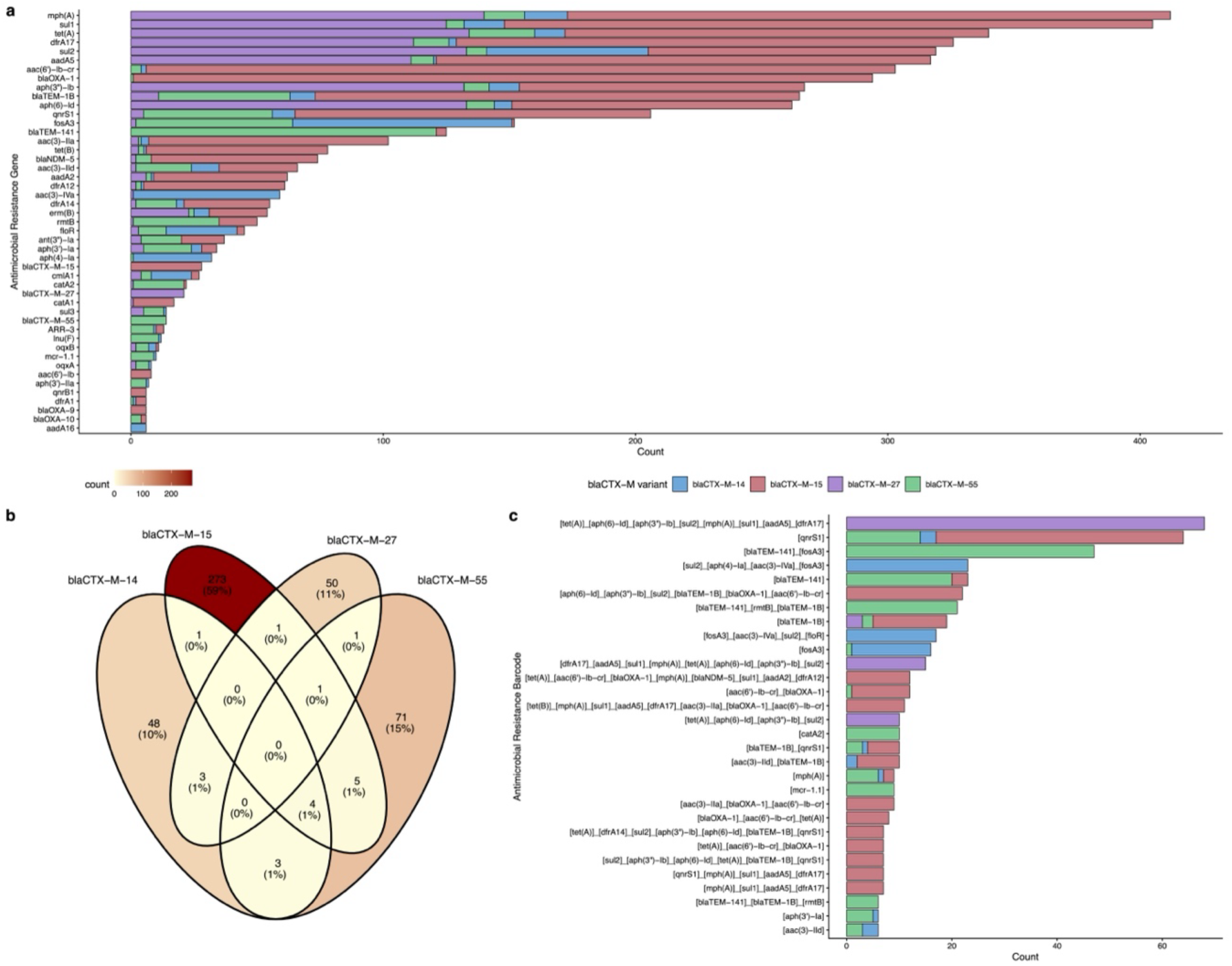
Distribution of antimicrobial resistance genes and resistance gene barcodes associated with the main plasmidic *bla*_CTX-M_ variants. **a,** Distribution of antimicrobial resistance genes (ARGs) identified within the variant-specific genomic windows surrounding plasmidic *bla*_CTX-M_ genes. Bars represent the number of occurrences of each resistance gene and are colored according to the associated *bla*_CTX-M_ variant (*bla*_CTX-M-14_, *bla*_CTX-M-15_, *bla*_CTX-M-27_, and *bla*_CTX-M-55_). Only ARGs detected more than 5 times are shown. **b,** Venn diagram showing the overlap of all 461 ARGs barcodes among the four main chromosomal *bla*_CTX-M_ variants. Each barcode corresponds to a unique combination of ARGs identified within the variant-specific genomic windows. Numbers indicate the count and percentage of barcodes belonging to each unique or shared category. The color gradient reflects the number of barcodes in each region. **c,** Distribution of ARG barcodes identified around plasmidic *bla*_CTX-M_ genes. Barcodes were defined as the ordered sequence of ARGs within genomic windows surrounding plasmidic *bla*_CTX-M_ genes and were displayed according to the orientation of the *bla*_CTX-M_ open reading frame. Bars represent the number of occurrences of each barcode and are colored according to the associated *bla*_CTX-M_. Only barcodes detected more than once are shown. All antimicrobial resistance genes barcodes are described in **Supplementary Table S2-d.**

**Extended Data Figure 7.**
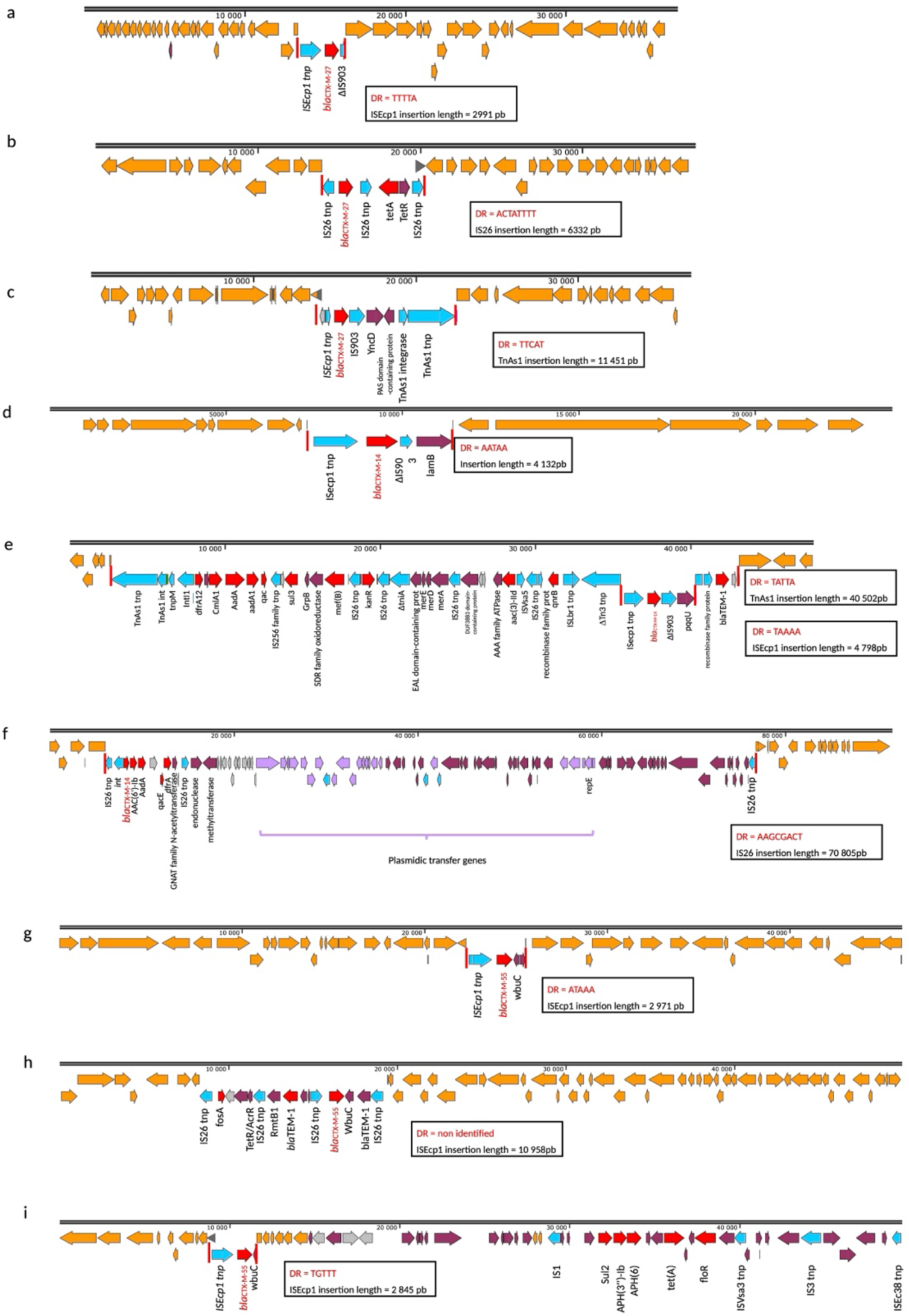
Chromosomal context of non-*bla*_CTX-M-15_ and mechanisms underlying their mobility on the chromosome. **a-c**, chromosomal context of *bla*_CTX-M-27_ (a:GCA_050198805.1; b:GCA_014169015.1, c:GCA_016820375); **d-f**, chromosomal context of *bla*_CTX-M-14_ (d:GCA_023658405.1; e:GCA_047782975.1, f: GCA_010724935.1); **g-i**, chromosomal context of *bla*_CTX-M-55_; (g:GCA_024271615.1 ; h: GCA_011492985.1, i: GCA_027944695.1). In orange, chromosomal *E. coli* genes; in red, antibiotic resistance genes; in blue, transposase genes; in grey, hypothetical protein coding genes; in purple, other coding genes. Red vertical bars indicate the target-site duplications.

**Extended Data Table 1.** IS element distribution across CTX-M variants and main resistance gene barcodes.

| CTX-M variant | Main barcode | IS associated |  |  |  |
| --- | --- | --- | --- | --- | --- |
|  |  | ISEcp1 (%) | IS26 (%) | Other (%) | Total |
| **CTX-M-15** | none | 189 (86.3) | 17 (7.8) | 13 (5.9) | **219** |
|  | *[aac(3)-IIa] [bla<sub>OXA-1</sub>] [aac(6')-Ib-cr]* | 1 (1.8) | 55 (98.2) | 0 (0.0) | **56** |
|  | *[qnrS1]* | 28 (90.3) | 2 (6.5) | 1 (3.2) | **31** |
|  | *[aac(3)-IIa] [bla<sub>OXA-1</sub>] [aac(6')-Ib-cr] [tet(A)]* | 19 (100.0) | 0 (0.0) | 0 (0.0) | **19** |
| **CTX-M-27** | none | 25 (75.8) | 5 (15.2) | 3 (9.1) | **33** |
|  | *[tet(A)]* | 0 (0.0) | 1 (50.0) | 1 (50.0) | **2** |
| **CTX-M-14** | none | 149 (94.9) | 4 (2.5) | 4 (2.5) | **157** |
|  | *[aac(3)-IId] [qnrS1] [bla<sub>TEM-1B</sub>]* | 5 (83.3) | 0 (0.0) | 1 (16.7) | **6** |
|  | *[aac(3)-IIa] [bla<sub>OXA-1</sub>] [aac(6')-Ib-cr]* | 2 (100.0) | 0 (0.0) | 0 (0.0) | **2** |
| **CTX-M-55** | none | 63 (95.5) | 2 (3.0) | 1 (1.5) | **66** |
|  | *[qnrS1]* | 3 (100.0) | 0 (0.0) | 0 (0.0) | **3** |

Values represent the number of chromosomal *bla*<sub>CTX-M</sub> insertions with the specified IS element (percentage of row total). Barcodes shown are the most frequent ARG combinations associated with each CTX-M variant. None: no additional co-integrated ARG. IS, insertion sequence.

## Notes

https://doi.org/10.5281/zenodo.20934804

### Competing Interest Statement

The authors have declared no competing interest.

