## Supplementary Figure S1 for "Chromosomal integration of *bla*_CTX-M_ redefines the evolutionary trajectory of antibiotic resistance in *E. coli* through lineage-structured expansion"

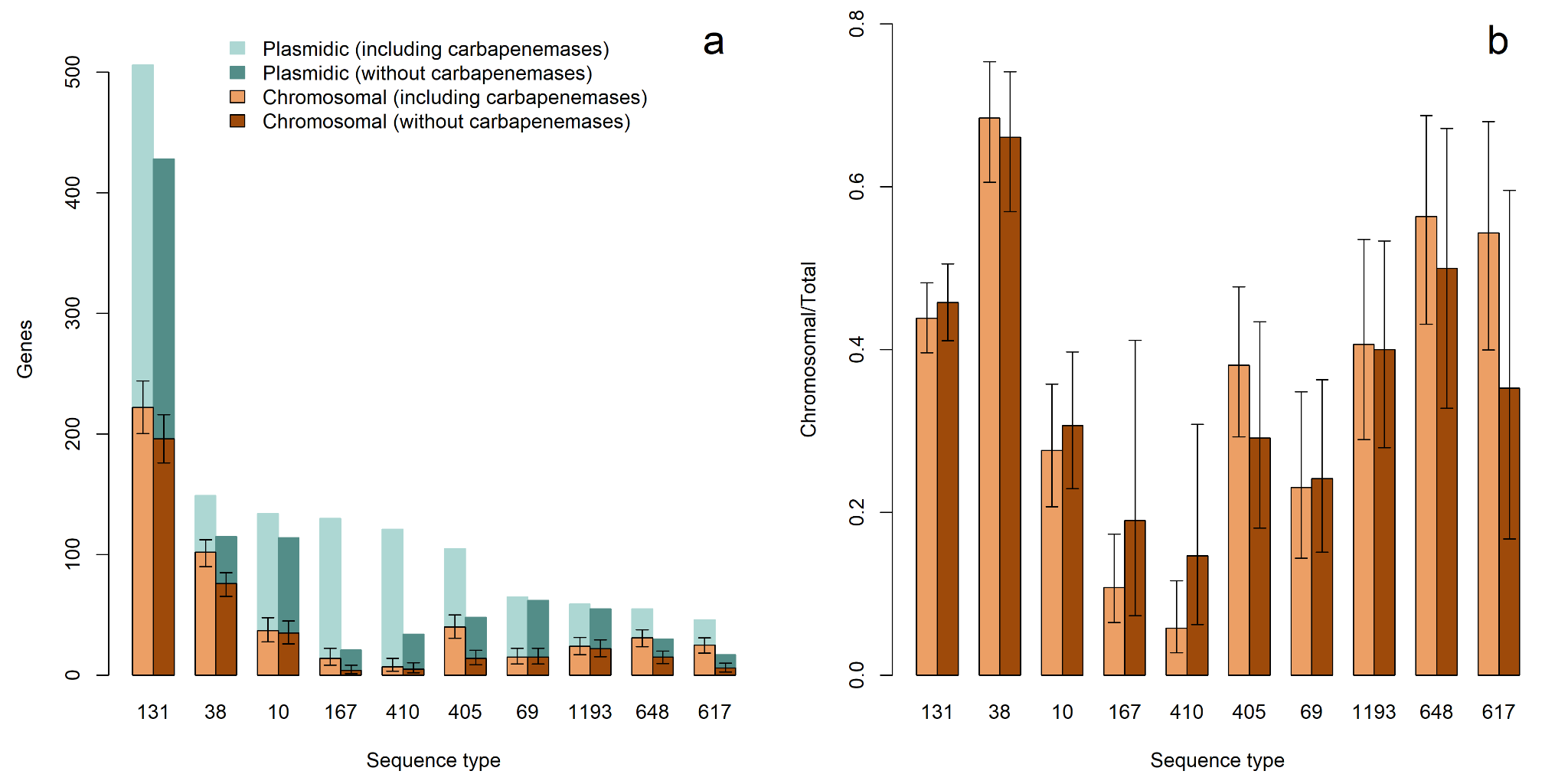

**Supplementary Figure S1. Interaction between chromosomal integration and the production of carbapenemases. a,** Number of *bla*_CTX-M_ genes assigned to plasmids (green) or to the chromosome (orange), considering the full dataset (light) with the dataset excluding carbapenemase-producing strains (dark). **b,** Corresponding proportion of *bla*_CTX-M_ genes assigned to the chromosome in each sequence type. **a-b,** Error bars indicate the 95% confidence intervals.
