## Supplementary Figure S2 for "Chromosomal integration of *bla*_CTX-M_ redefines the evolutionary trajectory of antibiotic resistance in *E. coli* through lineage-structured expansion"

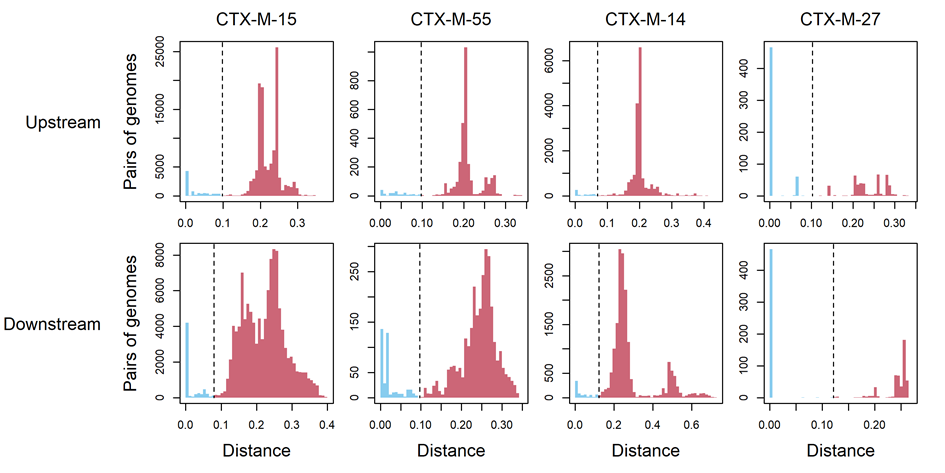

**Supplementary Figure S2. Clustering analysis to define unrelated pairs of genomes.** Distribution of the patristic distances between pairs of genomes for each variant (column) and direction (row), depending on whether they are related (blue) or unrelated (red). Each threshold (dashed vertical line) corresponds to the value separating the first cluster of distances according to the clustering analysis (blue) and the other clusters (red).
