## Supplementary Figure S3 for "Chromosomal integration of *bla*_CTX-M_ redefines the evolutionary trajectory of antibiotic resistance in *E. coli* through lineage-structured expansion"

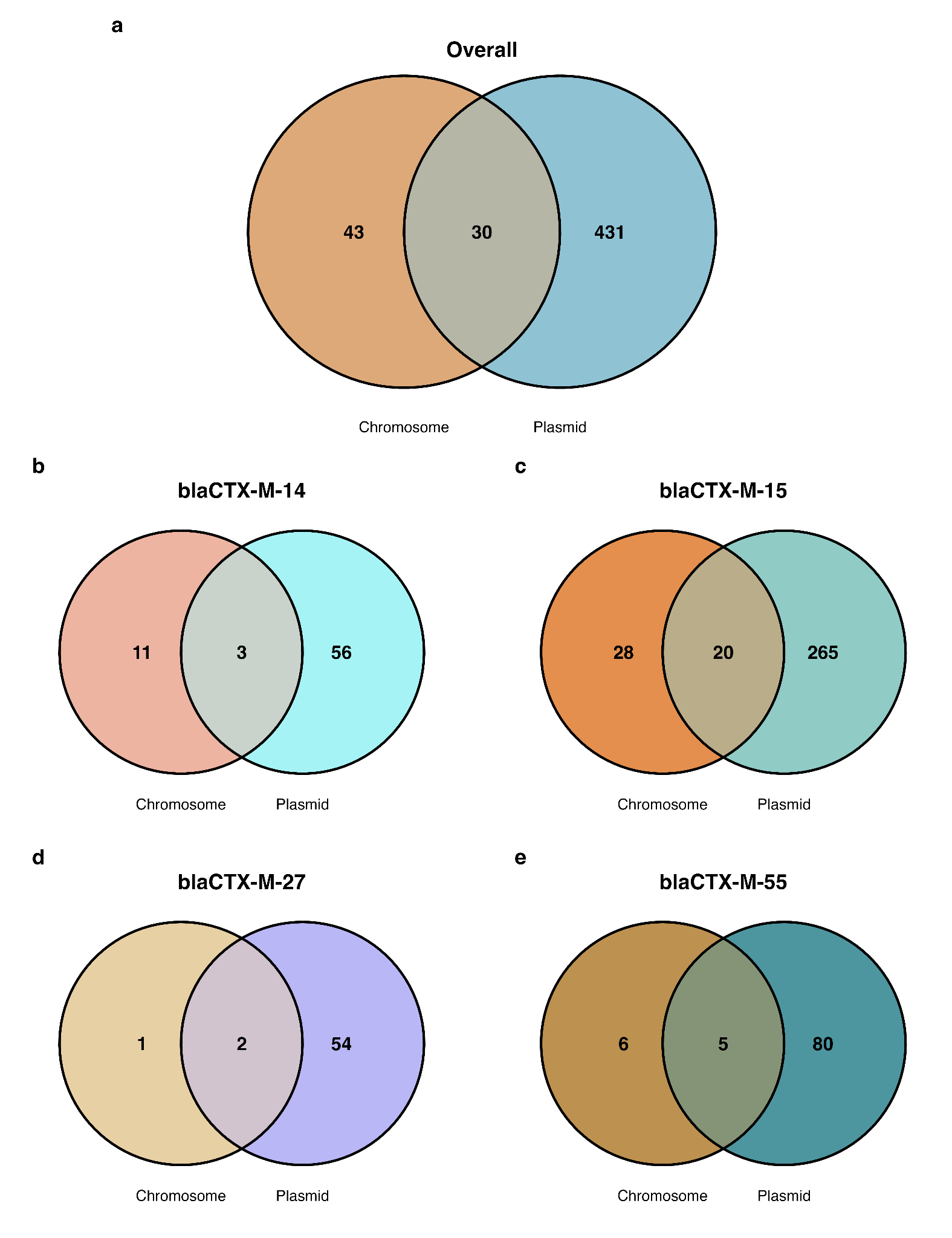

**Supplementary Figure S3. Distribution of ARG barcodes between chromosomal and plasmidic localization overall and across the main variants.**

Venn diagrams show the distribution of distinct ARG barcodes identified in chromosomal and plasmidic contexts. **a,** Overall distribution of ARG barcodes across the four major *bla*_CTX-M_ variants. **b-e,** Distribution of ARG barcodes for *bla*_CTX-M-14_, *bla*_CTX-M-15_, *bla*_CTX-M-27_, and *bla*_CTX-M-55_, respectively. Numbers indicate the number of distinct barcodes unique to chromosomal or plasmidic contexts, or shared between both localizations.
