## Supplementary Figure S4 for "Chromosomal integration of *bla*_CTX-M_ redefines the evolutionary trajectory of antibiotic resistance in *E. coli* through lineage-structured expansion"

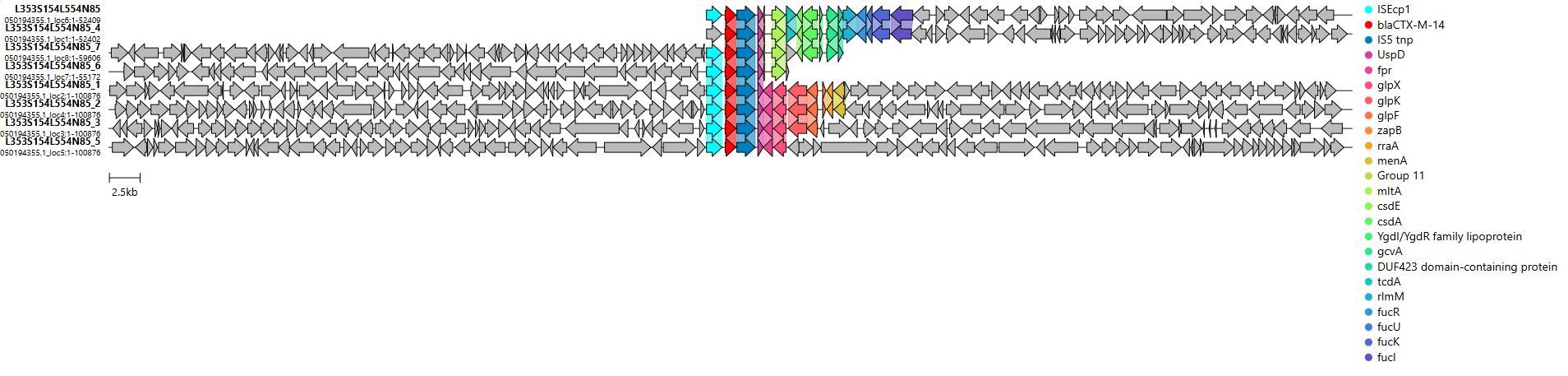

**Supplementary Figure S4. Synteny of the eight chromosomal *bla*_CTX-M-14_ integrations identified within a single strain.**

Each row represents one copy and its flanking genetic context (± 50 kb). Genes shared homology are coloured; chromosomal genes are shown in grey.
