## Supplementary Methods for "Chromosomal integration of *bla*_CTX-M_ redefines the evolutionary trajectory of antibiotic resistance in *E. coli* through lineage-structured expansion"

**Supplementary Materials and Methods**

Genomes were retrieved from NCBI, with metadata from EnteroBase. CTX-M variants were assigned to families using BLDB ^7^. Closely related genomes were removed based on cgMLST allelic distances ^8^, computed with chewBBACA v3.5.2 ^9^ using the *E. coli* Achtman cgMLST scheme ^10^. Phylogroups were assigned with ClermontTyper ^11^, O-serotypes with EcOH ^12^, *fimH* alleles with FimTyper ^13^, and insertion sequences with ISfinder ^14^. Pangenome analysis was performed with PPanGGOLiN v2.2.4 ^15^, and the phylogeny was inferred with FastTree v2.10.0 ^16^, and visualised with iTOL ^17^. Statistical analyses were performed in R ^18^. Mash-based trees were generated with Mashtree v1.4.6 ^3^.
