## Supplementary Note 1 for "Chromosomal integration of *bla*_CTX-M_ redefines the evolutionary trajectory of antibiotic resistance in *E. coli* through lineage-structured expansion"

**Estimation of the accumulation speed of chromosomal integration in the different subclades of ST131**

*Formulation of the model*

The following presents the methods of the model developed to approximate the dynamics of the chromosomal integration of *bla*_CTX-M_ genes in four subclades of ST131 (A, B, C1 and C2), from their last common ancestor to today.

Over this period, the variation in the proportion of chromosomal *bla*_CTX-M_ genes over time can reasonably be approximated by a sigmoidal function, under the assumption that the integration rate not only depends on the intrinsic characteristics of the strain, but also the frequency of *bla*_CTX-M_ genes themselves circulating in the bacterial population, which itself increased over time^1^. This approach results in a logit of the proportion of chromosomal *bla*_CTX-M_ genes increasing linearly over time. It is therefore akin to a logistical regression, consistent with the analysis comparing subclades presented in the main text of the article.

In this framework, the integration rate of subclade *i* at time *t* can be expressed as:

$$\begin{aligned} Y_{i}\left( t \right)=1/\left( 1+e^{-\left( logit\left( Y_{i}\left( t_{0} \right) \right)+r_{i}\left( t-t_{0} \right) \right)} \right),logit\left( y \right)=ln\left( y/\left( 1-y \right) \right),\#\left( 1 \right) \end{aligned}$$

with *t*_0_ the start of the time period considered, *Y_i_*(*t*_0_) the integration rate at the start of this period, and *r_i_* the accumulation speed of integration over this period (corresponding to the intrinsic rate of the sigmoidal function). For simplicity, the accumulation speed is assumed to be constant over a time period spanning from *t*_0_ to *t*_1_, such that *t*_0_ ≤ *t* < *t*_1_*.* Following Eq. 1, the value of *r_i_* can be expressed as:

$$\begin{aligned} r_{i}=logit\left( Y_{i}\left( t \right) \right)-logit\left( Y_{i}\left( t_{0} \right) \right)/\left( t-t_{0} \right).\#\left( 2 \right) \end{aligned}$$

The definition of periods and corresponding speeds for each subclade is based on their phylogenetic structure, which involves three main divergences (**Extended Data Fig. 4a**). The “A/B divergence” (separation of subclade B from subclade A) occurred circa 1890 ^2^, the “B/C” divergence (separation of subclade C from subclade B) circa 1990 ^2^, and the “C1/C2” divergence (separation of subclade C2 from subclade C1) circa 2000 ^2^. For model parsimony, a single accumulation speed is considered for each branch, with a single change resulting in a new speed at each divergence. Hence, four different accumulation speeds are defined: *r*_1_ is the ancestral speed prior to the A/B divergence, *r*_2_ results from a change in speed on the “B” branch after the A/B divergence, *r*_3_ results from a change in speed on the “C” branch after the B/C divergence, and *r*_4_ results from a change in speed on the “C2” branch after the C1/C2 divergence.

Therefore, the accumulation speed of subclade A is *r*_1_ from *t_AB_* (the date of the A/B divergence) to today, the accumulation speed of subclade B is *r*_2_ from *t_AB_* to today, the accumulation speed of subclade C1 is *r*_2_ from *t_AB_* to *t_BC_* (the date of the B/C divergence), then *r*_3_ from *t_BC_* to today, and the accumulation speed of subclade C2 is *r*_2_ from *t_AB_* to *t_BC_*, then *r*_3_ from *t_BC_* to *t_C1C2_* (the date of the C1/C2 divergence), then *r*_4_ from *t_C1C2_* to today (**Extended Data Fig. 4b**):

Based on this phylogenetic structure and Eq. 2, the four accumulation speeds can be expressed as:

$$\begin{aligned} \begin{matrix} \begin{matrix} r_{1}=\frac{logit\left( (Y_{A}(2025) \right) - logit\left( Y_{A}\left( t_{AB} \right) \right)}{2025 - t_{AB}} \\ r_{2}=\frac{logit\left( (Y_{B}(2025) \right) - logit\left( Y_{A}\left( t_{AB} \right) \right)}{2025 - t_{AB}} \end{matrix} \\ \begin{matrix} r_{3}=\frac{logit\left( (Y_{C1}(2025) \right) - logit\left( Y_{B}\left( t_{BC} \right) \right)}{2025 - t_{BC}},logit\left( Y_{B}\left( t_{BC} \right) \right)=logit\left( Y_{A}\left( t_{AB} \right) \right) +r_{2}\left( t_{BC}-t_{AB} \right) \\ r_{4}=\frac{logit\left( (Y_{C2}(2025) \right) - logit\left( Y_{C1}\left( t_{C1C2} \right) \right)}{2025 - t_{C1C2}},logit\left( Y_{C1}\left( t_{C1C2} \right) \right)=logit\left( Y_{A}\left( t_{AB} \right) \right) +r_{2}\left( t_{BC}-t_{AB} \right)+r_{3}\left( t_{C1C2}-t_{BC} \right) \end{matrix} \end{matrix}\#\left( 3 \right) \end{aligned}$$

Following Eq. 3, the accumulation speeds *r*_2_ through *r*_4_ can be expressed as differences from the previous speed, corresponding to the changes in speed during the A/B divergence (noted Δ*_AB_*), the B/C divergence (noted Δ*_BC_*) and the C1/C2 divergence (noted Δ*_C1C2_*):

$$\begin{aligned} \begin{matrix} r_{2}=r_{1}+\Delta_{AB},\Delta_{AB}=\frac{logit\left( (Y_{B}(2025) \right)-logit\left( (Y_{A}(2025) \right)}{2025-t_{AB}} \\ r_{3}=r_{2}+\Delta_{BC},\Delta_{BC}=\frac{logit\left( (Y_{C1}(2025) \right)-logit\left( Y_{B}\left( 2025 \right) \right)}{2025-t_{BC}} \\ r_{4}=r_{3}+\Delta_{C1C2},\Delta_{C1C2}=\frac{logit\left( (Y_{C2}(2025) \right)-logit\left( Y_{C1}\left( 2025 \right) \right)}{2025-t_{C1C2}} \end{matrix}\#\left( 4 \right) \end{aligned}$$

*Estimation of the accumulation speeds*

The values of *r*_1_, *r*_2_, *r*_3_, and *r*_4_ were numerically estimated together with their uncertainty, according to the model described above. To do so, distributions were defined for the underlying parameters, to reflect their uncertainty. The values of the current integration rate in the four subclades (*Y_A_*(2025), *Y_B_*(2025), *Y_C1_*(2025), and *Y_C2_*(2025)) were informed by the statistical analyses presented in the main text of this article. These analyses provided an estimate and standard error for each of them, which defined their respective distribution. The divergence dates (*t_AB_*, *t_BC_*, and *t_C1C2_*) were modelled as normal distributions, whose means were informed by the literature and whose standard deviations were set to 5 years. Hence, the 90% of the distributions of *t_AB_*, *t_BC_*, and *t_C1C2_* were respectively included the 1885 - 1890, 1985 - 1995, and 1995 - 2005 intervals. Finally, the integration at the time of the A/B divergence (*Y_A_*(*t_AB_*)) also had to be defined in order to estimate the four accumulation speeds, although its value had no impact on Δ*_AB_*, Δ*_BC_*, or Δ*_C1C2_*. Because *Y_i_*(*t*) > 0, as it is modelled by a logistic function, *Y_A_*(*t_AB_*) was set to an arbitrarily low value of 10^-12^. Following Eq. 3, multiplying (or dividing) *Y_A_*(*t_AB_*) by 1,000 decreases (or increases) all the speeds by 6.9. For the current integration of each subclade and each divergence date, a set of one million values was randomly drawn from their respective distributions, to compute one million values for each accumulation speed.

The estimated value of Δ*_AB_* (95%IC = [-0.009, 0.007]) was not significantly different from zero (**Extended Data Fig. 4c**), meaning that there was no significant difference between *r*_1_ and *r*_2_. However, Δ*_BC_* (95%IC = [-0.070, -0.005]) was significantly negative and Δ*_C1C2_* (95%IC = [0.039, 0.108]) was significantly positive, meaning that *r*_3_ was significantly smaller than *r*_2_ and *r*_4_. **Extended Data Fig. 4d** shows an estimated *r*_4_ generally higher than either *r*_1_ or *r*_2_.

Concerning the subclades themselves, our model therefore suggests that chromosomal integration of *bla*_CTX-M_ overall occurred at the same speed in subclades A and B, but that integration slowed markedly in subclade C following the B/C divergence. However, it also indicates that the integration accelerate significantly in subclade C2 after the C1/C2 divergence, to reach a higher integration rate than any other clade. The projected dynamics according to our model (for *Y_A_*(*t_AB_*) = 10^-12^) points towards a proportion of integrated *bla*_CTX-M_ in subclade C2 recently overtaking that of subclades A and B (**Extended Data Fig. 4e, 4f**).
