## Supplementary Note 2 for "Chromosomal integration of *bla*_CTX-M_ redefines the evolutionary trajectory of antibiotic resistance in *E. coli* through lineage-structured expansion"

**Definition of the window size considered for the study of synteny**

Studying synteny among *bla*_CTX-M_ insertions required defining the minimum size of the genomic region surrounding the gene to consider, in order to systematically encompass the entire inserted sequence. This minimum “window size” was defined here as the shortest sequence (in kilobases), upstream and downstream of the gene, for which pairs of sequences from unrelated genomes exhibited substantial divergence. Indeed, unrelated genomes carrying the same insertion were expected to display substantial greater differences in sequence outside the insert than within it.

*Identification of the unrelated pairs of genomes*

This approach first required the identification of pairs of unrelated genomes carrying the same *bla*_CTX-M_ variant. To achieve this, arbitrarily large regions either 100 kb upstream or 100 kb downstream of the gene were considered, based on the rationale that highly similar sequences over such large regions would indicate related genomes, regardless of the presence of the insertion. Corresponding mash-distances based phylogenetic trees were plotted using the Mashtree v1.4.6 software ^3^.

For each genome carrying one of the four main *bla*_CTX-M_ variants (CTX-M-15, CTX-M-55, CTX-M-14, CTX-M-27), the sequence either 100kb upstream or 100kb downstream of the gene was compared with those of all other genomes carrying the same variant. Patristic distances were computed for every pair of sequences for a given variant and a given direction (upstream or downstream). A clustering analysis was performed on each set of distances using a k-means algorithm. The value of *k* (number of clusters) was selected as the smallest value for which ratio of the between-cluster sum of squares to the total sum of squares exceeded 95% (**Supplementary** **Table S4**). In each analysis, the first cluster, ordered by increasing patristic distance, was assumed to contain the pairs of related sequences. The upper boundary of this cluster defined the threshold distance above which pairs of sequences were considered unrelated, i.e., distinguishable when considering a very large genomic region around the gene.

Pairs of unrelated genomes accounted for more than 90% of all possible pairs in every subset of a given variant and direction, except for sequences downstream of *bla*_CTX-M-55_ (87.1%) and for *bla*_CTX-M-27_ (54.5% upstream and 61.1% downstream). The latter was notably due to a large group of nearly identical genetic environments around the CTX-M-27 variant in ST131. The resulting threshold distances, which were used thereafter, ranged from 0.072 to 0.121 (**Supplementary Figure S2**).

*Definition of the minimal window size*

For each variant and direction (upstream or downstream), patristic distances between all pairs of unrelated genomes were computed over windows whose size ranged from 1kb to 100kb, with an increment of 1kb, starting from the gene. For a given pair of genomes, the minimum window size was defined as the smallest window for which the patristic distance exceeded the threshold defined above. The overall window size for a given variant and direction corresponded to the smallest size for which at least 95% of pairs of unrelated genomes exhibited a patristic distance greater than the corresponding threshold according to the clustering analysis (see above, **Supplementary Table S4**). Those resulting windows spanned the following intervals (in kb, with negative values corresponding to positions upstream of the gene): [-16, 33] for genomes with *bla*_CTX-M-15_, [-24, 5] for those with *bla*_CTX-M-55_, [-9, 14] for those with *bla*_CTX-M-14_, and [-15, 21] for those with *bla*_CTX-M-27_ (**Extended Data Fig. 5**). The position of the *bla*_CTX-M_ inside the window differed among variants. The gene was generally located closer to the beginning of the window in genomes with *bla*_CTX-M-15_ (**Extended Data Fig. 5a**), and closer to the end of the window for genomes with *bla*_CTX-M-15_ or *bla*_CTX-M-27_ (**Extended Data Fig. 5b and d**).

*Characterization of the genetic background of chromosomally integrated variants*

We used the defined windows to explore the genetic background surrounding the chromosomally integrated *bla*_CTX-M_ genes. Thus, we sought to characterize the barcode content of the genes, which is comprised of sequential combinations of Antimicrobial Resistance Genes (ARGs) upstream and downstream, and the synteny among the genetic environment.
