## Supplementary Note 3 for "Chromosomal integration of *bla*_CTX-M_ redefines the evolutionary trajectory of antibiotic resistance in *E. coli* through lineage-structured expansion"

**Characterization of *bla*_CTX-M_ chromosomal insertion events**

For each *bla*_CTX-M_ variant, the genomic windows flanking the gene, identified during the localization step described above, were extracted and aligned by synteny analysis ^4^ to search for conserved sequence motifs. Identification of insertion sequences shared across multiple strains revealed two predominant insertion sequences potentially implicated in gene integration, ISEcp1 and IS26. A subset of insertions, corresponding to the most frequent events, was manually re-annotated using SnapGene (SnapGene software, Dotmatics), ISfinder ^5^ and TnCentral ^6^ to precisely determine their insertion length.

**Definition of the mechanism of mobility**

Each chromosomal *bla*_CTX-M_ insertion was assigned to one of three categories based on suspected mobility mechanism:

- **ISEcp1-associated**: defined by the presence of a complete ISEcp1 transposase sequence located immediately upstream of *bla*_CTX-M_ identified within the flanking *Escherichia coli* chromosomal gene bracketing the insertion site identified by clinker or more precisely on SnapGene with 5-bp target site duplication (TSD).
- **IS26-associated**: defined by the presence of at least two complete IS26 transposase sequences flanking *bla*_CTX-M_ on either side or more precisely on SnapGene.
- **Other**: this category pools insertions attributed to alternative, less frequent resistance-mobilization mechanisms (Tn2 family…) and with those for which the underlying mechanism could not be identified.
