## Supplementary Table S1 for "Chromosomal integration of *bla*_CTX-M_ redefines the evolutionary trajectory of antibiotic resistance in *E. coli* through lineage-structured expansion"

**Supplementary Table S1. Parameter estimates of the logistic regression on sequence type integration rates**

| Factor | Estimate (logit) | Standard error | p-value |
| --- | --- | --- | --- |
| Intercept (ST131) | -0.246 | 0.090 | 0.006 |
| **ST410** | **-2.544** | **0.400** | **<0.001** |
| **ST167** | **-1.868** | **0.297** | **<0.001** |
| **ST69** | **-0.958** | **0.308** | **0.002** |
| **ST10** | **-0.717** | **0.213** | **0.001** |
| ST405 | -0.239 | 0.220 | 0.277 |
| ST1193 | -0.131 | 0.280 | 0.640 |
| ST617 | 0.421 | 0.309 | 0.174 |
| ST648 | 0.502 | 0.286 | 0.079 |
| **ST38** | **1.021** | **0.198** | **<0.001** |

Parameter estimates of the logistic regression on sequence type integration rates are presented with their standard error and p-value. The rate of ST131 is used as reference and other parameters account for differences with ST 131. The ST showing a significant difference with ST131 are in bold.
