## Supplementary Table S3 for "Chromosomal integration of *bla*_CTX-M_ redefines the evolutionary trajectory of antibiotic resistance in *E. coli* through lineage-structured expansion"

**Supplementary Table S3. Parameter estimates of the logistic regression on ST131 clade integration rates**

| Factor | Estimate (logit) | Standard error | p-value |
| --- | --- | --- | --- |
| Intercept (clade C1) | -1.197 | 0.176 | <0.001 |
| **Clade A** | **1.383** | **0.279** | **<0.001** |
| **Clades B** | **1.197** | **0.530** | **0.024** |
| **Clade C2** | **1.550** | **0.226** | **<0.001** |

Parameter estimates of the logistic regression on ST131 clade integration rates are presented with their standard error and p-value. The rate of clade C1 is used as reference and other parameters account for differences with clade C1. The ST showing a significant difference with clade C1 are in bold.
