## Supplementary Table S4 for "Chromosomal integration of *bla*_CTX-M_ redefines the evolutionary trajectory of antibiotic resistance in *E. coli* through lineage-structured expansion"

**Supplementary Table S4. Results from the clustering analyses using the k-means algorithm for each variant and direction.**

| Variant | Direction | Number of clusters | Ratio of square sums | Threshold distance |
| --- | --- | --- | --- | --- |
| CTX-M-15 | upstream | 5 | 0.957 | 0.100 |
| CTX-M-15 | downstream | 6 | 0.956 | 0.079 |
| CTX-M-55 | upstream | 4 | 0.950 | 0.098 |
| CTX-M-55 | downstream | 5 | 0.964 | 0.097 |
| CTX-M-14 | upstream | 7 | 0.956 | 0.072 |
| CTX-M-14 | downstream | 6 | 0.968 | 0.105 |
| CTX-M-27 | upstream | 3 | 0.962 | 0.092 |
| CTX-M-27 | downstream | 2 | 0.964 | 0.121 |

The number of clusters used, the ratio of the between-cluster sum of squares to the total sum of squares, as well as the threshold distance are indicated for each analysis.
